# Re-activation of neurogenic niches in aging brain

**DOI:** 10.1101/2024.01.27.575940

**Authors:** Roy Maimon, Carlos Chillon-Marinas, Sonia Vazquez-Sanchez, Colin Kern, Kresna Jenie, Kseniya Malukhina, Stephen Moore, Jess Cui, Alexander Goginashvili, Siavash Moghadami, Alexander Monell, Melissa McAlonis-Downes, Christine Hong, Paymaan Jafar-Nejad, C. Frank Bennett, Quan Zhu, John Ravits, Don W. Cleveland, Bogdan Bintu

## Abstract

Recent studies proposing induced glia-to-neuron conversion raised the potential for generating new neurons to replace those lost due to injury, aging or neurodegenerative diseases. Here, single-cell spatial transcriptomics [<u>M</u>ultiplexed <u>E</u>rror <u>R</u>obust <u>F</u>luorescence *In <u>S</u>itu* <u>H</u>ybridization (MERFISH)] is used to construct a spatial cell atlas of the subventricular and dentate gyrus neurogenic niches of young and aged adult murine brain. RNAs that encode the RNA binding protein <u>P</u>olypyrimidine <u>T</u>ract-<u>B</u>inding <u>P</u>rotein (PTBP1) in the aged murine brain are determined to be highest in glia that line previously active neurogenic niches. A glial cell population with ependymal character within an initially quiescent subventricular neurogenic niche in the aged murine brain is identified that upon transient suppression of PTBP1 reenters the cell cycle, replicates DNA, and converts into neurons through a canonical adult neurogenesis pathway. Glia-derived neurons migrate from this niche, with some neurons transiting to the striatum and acquiring a transcriptome characteristic of GABAergic inhibitory neurons. Similar PTBP1 expressing quiescent glia are identified in the corresponding neurogenic niche of aged human brain. Thus, transient reduction of PTBP1 holds potential for inducing the generation of new neurons in quiescent neurogenic niches of the aged nervous system, thereby offering promising therapeutic applications.

**Bullet point summary:**

1) Single-cell spatial transcriptomics is used to validate active neurogenesis in the two neurogenic niches of the young adult murine brain, determine that those niches are quiescent in the aging adult brain of mice, and demonstrate the absence of neurogenesis in the aging human brain.
2) The RNA binding protein PTBP1 is determined to be most highly expressed within glia that line the aged murine and human neurogenic niches, with its transient reduction sufficient in mice to activate/re-activate expression of genes characteristic of immature neurons.
3) Suppression of PTBP1 using a single intra-cerebral-ventricular injection of PTBP1–targeting antisense oligonucleotide (ASO) induces generation of new immature neurons in the neurogenic niches of the aged mouse brain via a canonical adult neurogenesis pathway.
4) Single-cell RNA signature tracing is used to identify *a)* a subclass of ependymal cells in a previously quiescent neurogenic niche of the aged mouse brain that convert into GABAergic inhibitory neurons following transient suppression of PTBP1, and *b)* the molecular steps in the conversion process including cell cycle re-entry, DNA replication, and transcriptome changes that mimic canonical neurogenesis.
5) A similar class of PTBP1-expressing ependymal cells lining the ventricle of the aging non-human primate and human brains is identified, suggesting the promise of re-activation of neurogenesis as a therapeutic approach in humans.

## Introduction

Since neuronal populations are mostly determined during embryogenesis, loss of neurons in the adult has been thought to be irreversible, with no replacement of those lost due to neurodegeneration, injury, or aging. Indeed, more than 100 years ago, Ramon y Cajal stated^1^: “*In the adult centers, the neural paths are something fixed and immutable: everything may die, nothing may be regenerated”*, establishing a central dogma for neuroscience during the last century. Cajal also directly challenged the neuroscience community to identify ways to overcome this barrier: “*It is a duty for future generations find a way to overcome the intrinsic failure of adult brain to regenerate*”. Intuitively, replacing damaged or dying neurons with newly generated ones would be an approach to reverse disease or age-dependent damage.

In 1987, the pioneering work of Weintraub and colleagues first demonstrated that a fibroblast could be directly converted into a muscle cell by forced expression of the MyoD transcription factor^2,3^. Conversion of cells in the context of the brain was achieved twenty-five years later when Gotz and her team^4^ reported the concept of glia-to-neuron conversion by chronic expression of a neuronal transcription facto Pax6. Since then, the development of methods to reprogram somatic human cells into induced pluripotent stem cells (iPSCs) enabled the generation *in vitro* of a wide range of neural cells from any human individual^5^. Although such induced neurons are now widely used experimentally, the appropriateness of iPSC-derived neurons therapeutically has been questioned by many investigators, since this approach can lead to detrimental outcomes including tumorigenesis^6^, genetic aberrations^7^, and reset of epigenetic and aging signatures^8^. Recognizing this, direct *in vivo* conversion into neurons of non-dividing cells expressing an individual’s own histocompatibility antigens might be a more attractive approach for therapeutic uses and could establish a new approach for treating neuronal loss.

Several groups have reported success in astrocyte-to-neuron conversion in culture^9–15^ and initiated testing this concept *in vivo*, in the mammalian brain. Key transcription factors that have been reported to induce astrocyte-into-neuron conversion, alone or in combination with other factors, in mouse brain include: each of four bhlh transcription factors (Neurog2^16–22^, Ascl1^23–32^, Atoh(s)^33^, and NeuroD1^34–37^), a homeobox protein Pax6^38^, Sox2^39–43^, Dlx2^31,44^, and Oct4^45^. Inhibition of the RNA binding protein PTBP1 was also reported to facilitate generation of new neurons in the brain^46–50^, including a report that functional, replacement of nigral neurons could be induced by PTBP1 reduction within GFAP-labelled glia after unilateral injection of a toxin to kill most nigral neurons, thereby producing chemically-induced Parkinson’s-like disease in mice^51^. However, in 2021 prominent challenges arose^52–54^ (reviewed by Berninger^55^). First, Zhang and colleagues reported that the apparent new neurons produced through AAV delivery of transcription factors or shRNA suppression of PTBP1 were actually mistakenly labelled pre-existing, morphologically mature neurons^52^, rather than converting from astrocytes into neurons. Second, Blackshaw and colleagues reported that inducible genetic disruption of PTBP1 did not generate new neurons from mature astrocytes in normal adult mouse retina^53^ or brain^56^. While other groups made claims to reinforce these challenges^54,57^, additional evidence was reported in support of PTBP1 suppression in generating new neurons^58^. Using a therapeutically feasible approach^59,60^ with an antisense oligonucleotide (ASO) delivered by injection into cerebral spinal fluid (CSF), we reported that transient suppression of PTBP1 was sufficient to trigger quiescent radial glial-like cells in the mouse dentate gyrus to convert into new neurons in aged adult mice (an age ≥7 months after the last documented “adult” neurogenesis^61,62^) through a process involving transient induction of the early neuronal marker doublecortin (Dcx), functional integration into endogenous circuits, and enhancement of memory performance^46^.

Single-cell and single-nucleus transcriptomic methods^63–65^ are rapidly changing our understanding of the cellular composition and role of the different cell types within the mammalian brain^63–69^, with hundreds of transcriptionally distinct cell populations already identified and characterized^65,69^. While these methods provided important insights into the pathways of generation of new neurons in the developing and adult brain^69–71^, preservation of spatial information and reliable quantification of expression is key to analysis of these rare events, which take place within a complex cellular environment. Here we used a spatial transcriptomics method with a high degree of spatial resolution and detection efficiency called MERFISH^63–65,72,73^ (<u>M</u>ultiplexed <u>E</u>rror <u>R</u>obust <u>F</u>luorescence *In <u>S</u>itu* <u>H</u>ybridization). Unlike current “conventional” single cell (or single nucleus) RNA sequencing methods, this approach can quantify more accurately genes expressed in individual cells/nuclei and can retain the spatial context and morphology of cells within the tissue. We couple MERFISH with transient, ASO-mediated suppression of PTBP1 to 1) identify a glial population with ependymal cell character located within an initially quiescent subventricular neurogenic niche in the aged murine brain that converts into new neurons and 2) determine that conversion follows a canonical adult neurogenesis pathway including cell division and proliferation.

## Results

### A cell atlas of the neurogenic niches of the aging mouse brain

We utilized a MERFISH spatial transcriptomics, a method previously used to produce a spatial atlas of murine brain hippothalamus^66^, primary motor cortex^67^, cerebellum, and other brain regions including a recent profiling of the near entire mouse brain^65,74–76^. In this approach, up to a thousand genes are first targeted and hybridized with an average of 50 single-molecule FISH probes per gene. Using error-robust barcoding, combinatorial labeling, and sequential imaging, RNAs are encoded with unique N-bit binary words through hybridization with unlabeled encoding probes. The RNA molecules are then labelled and imaged combinatorically across multiple rounds of hybridization. The RNA with binary word “101…1” is labeled with encoding probes that contain the 1st, 3rd, … Nth readout sequences and are detected with corresponding readout probes. This approach enables hundreds of distinct genes to be imaged and their expression quantified in only 10-20 rounds of imaging ***(Figure 1A)***.

**Figure 1:**
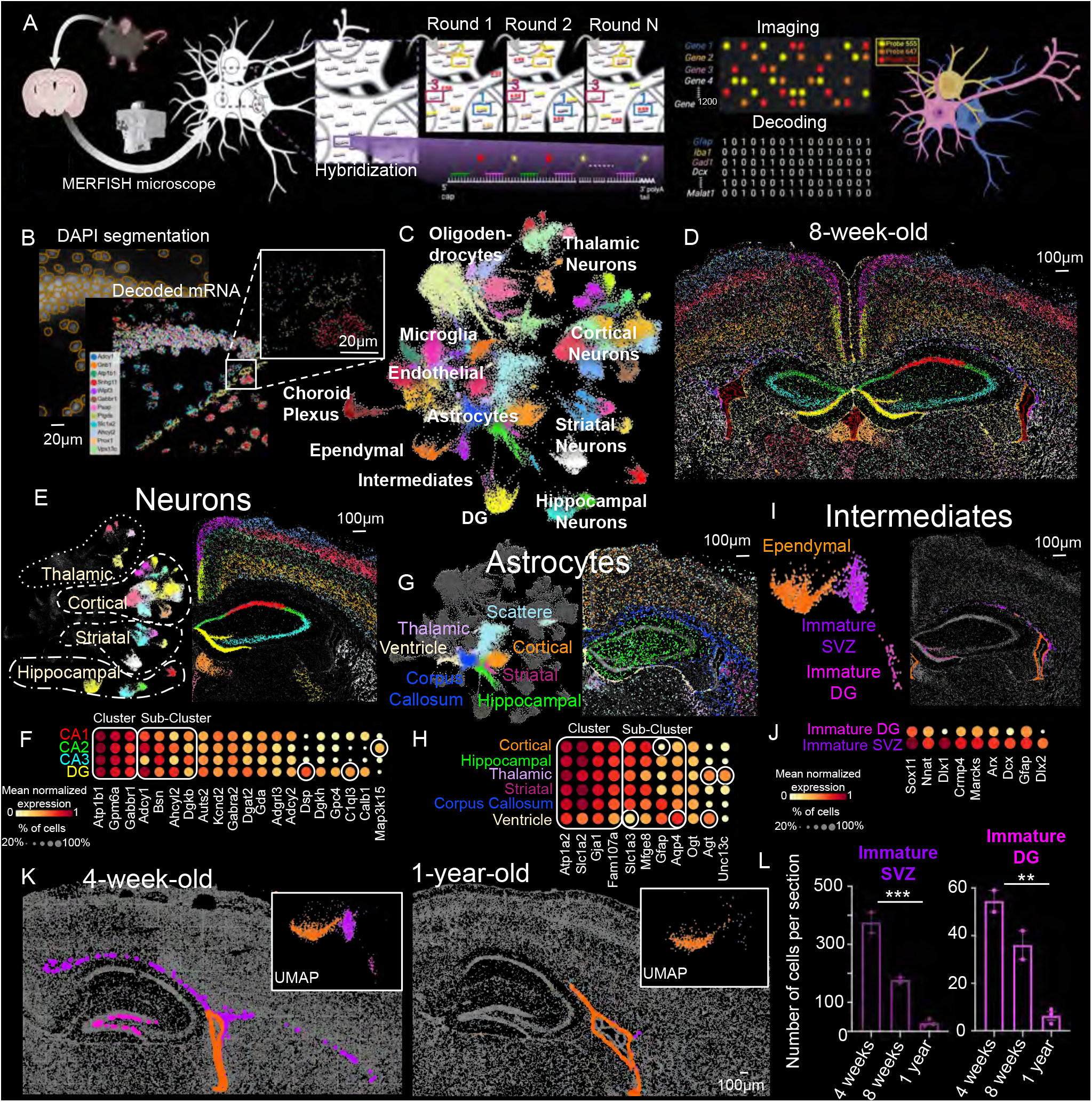
The spatial brain cell atlas of adult neurogenesis in young and aged mice. **(A)** Scheme depicting MERFISH imaging. Sagittal or coronal 16μm-thick brain sections were hybridized with a primary library (∼50 oligonucleotide probes per mRNA species) targeting 223-273 different mRNAs. A microfluidics enabled microscope was used to hybridize and image (2-3 colors across 11-16 cycles) the transcripts of mRNAs in each single cell. Combinatorial and error robust imaging strategies were employed to allow precise identification of the transcripts^63^. **(B)** A representative image of the dentate gyrus of an 8-week-old mouse brain, with the transcripts of 12 genes depicted with different colors (Adcy1 (blue), Gnb1 (orange), Atp1b1 (green), Snhg11 (red), Wipf3 (purple), Gabbr1 (brown), Psap (pink), Ptgds (green), Slc1a2 (light blue), Ahcyl2 (light purple), Prox1 (light orange), Vps13c (light green)). Left panel: Individual cells were segmented by a machine learning algorithm (Cellpose^86^) based on PolyA and DAPI stainings. **(C)** UMAP representation of ∼80k cells and 223 genes. 67 cell types were identified via Leiden clustering and manually annotated. **(D)** Spatial map of 8-week-old mouse brain crossection showing the cell types from (C) with matching colors. **(E,G,I)** UMAPs and spatial maps of the subclasses of **(E)** hippocampal neurons (4 clusters), **(G)** astrocytes (8 clusters), and **(I)** intermediate cell types (3 clusters)**. (F,H,J)** Dot plots representing the most differentially expressed genes in **(F)** hippocampal neurons, **(H)** astrocytes, and **(J)** intermediates cells. White squares mark the top genes defining the clusters. White circles highlight genes previously reported as corresponding cell type markers^90^. **(K)** Representative UMAP and spatial maps of intermediate/immature cells comparing 4-weeks-old mouse sagittal brain section (left) with aged 1-year-old brain (right). Orange: Ependymal cells, Purple: SVZ intermediates, Pink: DG intermediates. **(L)** Quantification of the number of immature cells in 4-weeks-old, 8-weeks-old, and 1-year-old mouse brains (n□=□2,2,3 independent brain sections, respectively; One-way ANOVA, **p□=□0.0035, ***p□=□0.0003).

We implemented MERFISH with a gene list focused on determining the cell identities of neurons, glia, and cell intermediates appearing during canonical adult neurogenesis^71,77^. We selected a total of 223 genes ***(Figure S1A,B)*** including genes covering multiple transcription factor classes^78^ (*Pax6, Sox2*, etc.), genes expressed within cell types of neurogenic niches^79,80^ (e.g., *Aqp4* and *Glast* marking astrocytes, *Ascl1* and *Hes5* marking radial glial like cells, *Dcx* and *NeuroD1* marking immature neurons, *Rbfox3*, *Calb1* and *Prox1* marking mature dental gyrus neurons, etc.), genes covering axonal guidance and cell adhesion molecules^81^ (i.e., *Rubo2*, *Sema3A*, *BDNF* etc.), genes covering ion channels and neurotransmitters (*Gabra1*, *Gabra2*), and genes within or related to a proposed PTBP1 pathway^82–84^ (e.g., *PTBP1, PTBP2, BRN2, REST, CLASP1*). An additional set of 110 genes was selected based on the Allen Institute’s single-nuclei RNA-sequencing (snRNA-seq) dataset^85^ to capture finer cell type divisions within hippocampal and cortical neurons and their glial partners (***see Methods***).

MERFISH imaging of these 223 genes was first performed in 16-μm thick coronal sections of the young (8-week-old) adult murine brain. Nuclear staining (DAPI), coupled with the Cellpose machine learning approach^86^, were performed to enable cell segmentation and associate each mRNA transcript with its corresponding cell ***(Figure 1B)***. Based on the RNA contents of each single-cell imaged, we performed unsupervised clustering (using the Leiden algorithm^87^) and identified 59 cell type clusters from the 83,981 cell expression profiles within a single coronal section ***(Figure 1C)***. We first annotated each of these clusters into the major cell-type classes based on the expression of the expected marker genes. These included cortical layers L2, L2/3, L3, L5 and L6 neurons, hippocampal neurons, thalamic and striatal neurons, astrocytes, oligodendrocytes and their precursors, microglia, ependymal, and epithelial cells. Good correlation of the RNA expression profiles was found between the MERFISH-defined cell types and the corresponding cell-type classifications from a recently published single-cell sequencing cell atlas of the murine brain^88^ ***(Figure S1C,D,E)***.

Furthermore, while defining the 59 cell types solely on the RNA contents of each cell, each cell type was identified to have a distinct spatial organization within the section ***(Figure 1D)***. For instance, the different subtypes of neurons displayed the expected layered organization within the cortex: from the outer layer 2 to the inner layer 6 ***(Figure 1E)***. The hippocampal formation was divided into its expected anatomy with the three Cornu Ammonis subdivisions CA1, CA2, CA3 superior to the Dentate Gyrus (DG) neurons with unique gene expression corresponding to each cluster ***(Figure 1E)***. For instance, within hippocampal neurons, the *Dsp* gene encoding desmoplakin, a protein which functions to maintain the structural integrity of cell contacts^89^, was found highly expressed only within the layer of GABAergic inhibitory granular cell neurons of the DG ***(Figure 1F)***. Seven different classes of astrocytes were identified, each with a highly distinctive spatial organization ***(Figure 1G)***. Six of the 7 groups were almost exclusively localized within cortical, hippocampal, striatal, thalamic, ventricular, and corpus callusom brain regions, respectively (**Figure 1G**). In contrast, the seventh astrocyte type was diffusely positioned across all major brain regions. While all astrocyte classes expressed global astrocytic markers (e.g., *Gja1* and *Slc1a2*), each different subtype was distinguished by expression of specific gene combinations. For instance, the combination of *Agt*, *Gfap*, and *Slc1a3* expression was selectively enriched in ventricular astrocytes whereas *Unc13c* expression was enriched in thalamic and cortical astrocytes (**Figure 1H**), findings consistent with a prior report using single molecule FISH (smFISH) to define five subclasses of astrocytes^90^.

Among the cell-type clusters identified within this initial MERFISH data, two small populations of cells (229 subventricular immature cells (purple) and 23 subgranular dentate gyrus immature cells (pink) were characterized by high expression of neurogenic genes (e.g., *Nnat, Dcx, Dlx1,2* and *Sox11*) ***(Figure 1I,J)***. Based on the physical location of these cells and the genes that they expressed within the young adult brain, we hypothesized that these populations represent intermediate states of adult neurogenesis within the subventricular zone (SVZ) and the subgranular zone of the dentate gyrus (SGZ)^79,80^.

We enriched our MERFISH library to include 52 additional neurogenic gene markers (e.g., *Igfbpl1, Sox4*, etc) (***Figure S1F)*** selected based on recent single-cell sequencing data of the SVZ^71^ and the SGZ^70^. We performed MERFISH across mice of different ages including younger (4-week-old) and aged (1-year-old) brains ***(Figure 1K,)***. The number of immature cells was higher in young brains ***(Figure 1L)*** within the subventricular and subgranular zones, with an average of 375 immature SVZ cells and 54 immature SGZ cells found within single images containing an average of 80,000 cells. Neurogenesis across multiple 1-year-old brains was highly reduced ***(Figure 1L)***, with only an average of 29 and 6 immature neurons in comparable images within SVZ and SGZ, respectively.

### PTBP1 is highly expressed within a subpopulation of glial cells in the SVZ

Recognizing that multiple prior efforts have proposed that the gene encoding PTBP1 (denoted *PTBP1* in humans and *Ptbp1* in mice) is an essential developmental gene involved in the neurogenesis pathway^82,91^, we utilized MERFISH to quantify the remaining PTBP1 expression within each cell type of the aged (1-year-old) mouse brain ***(Figure 2A)***. *Ptbp1* RNAs were highest in two cell populations in the SVZ: ependymal cells within the choroid plexus and the lateral ventricle wall ***(Figure 2A, B, C)*** Consistent with this, PTBP1 protein (measured using immunofluorescence imaging) was highest in these two areas ***(Figure 2D)***. *Ptbp1* mRNAs accumulated to modest levels in the seven astrocytic classes identified by MERFISH ***(Figure 2C,E)*** and in neurons within the CA1, L2/3, L2-3, DG, CA3 regions of the hippocampus and cortex ***(Figure 2C,F)***. Proteins whose expression is expected to depend on PTBP1^82,91,92^ ***(Figure 2G)*** include *Rest* (also known as NRSF [Neuron-Restrictive Silencer Factor]), a repressive transcription factor whose function has been proposed to inhibit the expression of neuronal genes in non-neuronal cells^93,94^. Consistent with this, levels of mRNAs encoding Rest correlated with *Ptbp1* mRNAs, with the highest levels within choroid plexus and ventricular ependymal cells and lower expression in astrocytic and neuronal clusters ***(Figure 2H)***. RNAs encoding *Ptbp2,* another member PTB family member, were negatively corelated with those encoding *Ptbp1*, with highest expression within L3 neuronal cell types and lowest within ependymal cells ***(Figure 2I)***, consistent with previous proposals that these two members of the PTB family inhibit each other^83^. Genes previously implicated to be upregulated (*Rtn4*) or downregulated (*Brn2*, *Clasp1*, and *NeuroD1*) by PTBP1 action were found in our MERFISH imaging to correlate/anticorrelate (as predicted) with *Ptbp1* RNA levels in multiple classes of neurons and astrocytes ***(Figure 2J)***.

**Figure 2:**
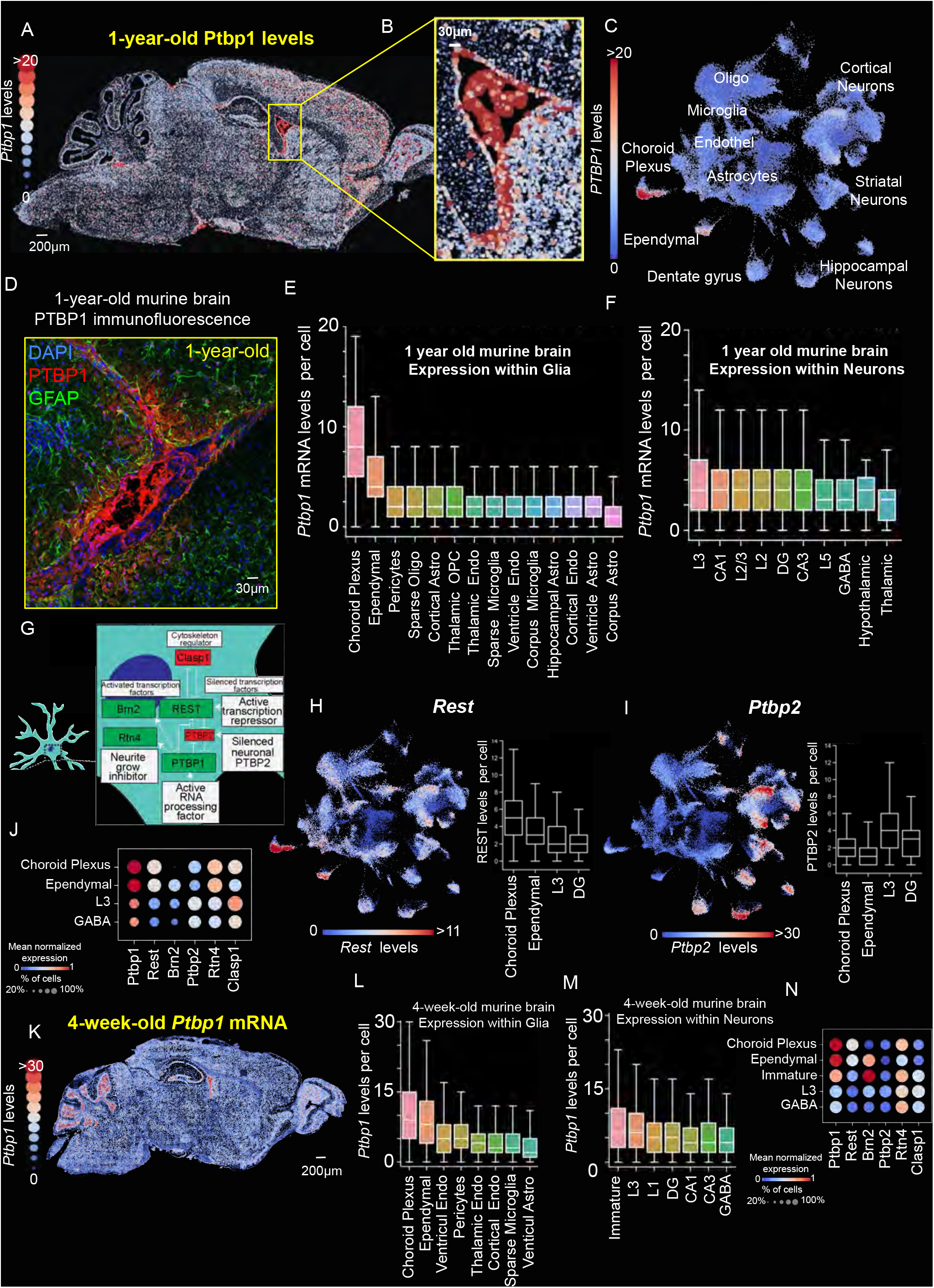
PTBP1 is highly expressed within a subtype of glia in the adult neurogenic niches. **(A)** Representative image showing *Ptbp1* mRNA expression levels with single-cell resolution in a sagittal section of a wild type 1-year-old murine brain. **(B)** Inset of (A) showing the ventricular zone. **(C)** UMAP showing single-cell *Ptbp1* mRNA expression in a 1-year-old mouse brain. The coordinates of the UMAP from Figure 1C were used for consistency. **(D)** Representative immunofluorescence image of the ventricular zone of a 1-year-old murine brain. PTBP1 (Red), GFAP (green), and DAPI (blue). **(E, F)** Box plots showing the *Ptbp1* mRNA expression levels per cell in glial **(E)** and neuronal **(F)** subtypes. **(G)** Schematic of the proposed PTBP1 regulatory loop^82,92^ including genes: *Ptbp2*, *Rest*, *Brn2*, *Rtn4* and *Clasp1*. **(H)** Left: UMAP quantifying *Rest* single-cell mRNA levels in a 1-year-old mouse brain. Right: Box plots showing *Rest* mRNA expression levels per cell in representative neuronal and glial cell types. **(I)** Same as (H) for *Ptbp2*. **(J)** Dot plot representing the mRNA levels per cell of *Ptbp1* and proposed PTBP1-controlled genes (*Rest*, *Brn2, Ptbp2, Rtn4, Clasp1*) in Choroid plexus cells, Ependymal cells, L3 cortical neurons and DG neurons. **(K)** Representative brain projection map of *Ptbp1* mRNA expression in a sagittal section of a 4-week-old murine brain. **(L,M)** Box plots quantifying the *Ptbp1* mRNA levels per cell in glial **(L)** and neuronal **(M)** populations of a 4-week-old murine brain. **(N)** Dot plot representing the mRNA levels per cell of proposed PTBP1 dependent genes (*Rest*, *Brn2, Ptbp2, Rtn4, Clasp1*) in Choroid plexus cells, Ependymal cells, immature neurons, L3 cortical neurons and DG neurons.

Consistent with what we found in 1 year old mice, in the brains of young adult (4 week old) mice in which active neurogenesis was ongoing, *Ptbp1* RNA ***(Figure 2K,,M)*** levels were highest in ependymal cells of the choroid plexus and SVZ, with intermediate levels in immature neuronal-like cells, and only low levels in mature glia ***(Figure 2K,L,M)***. These results are consistent with a current model of a canonical neurodevelopment in which PTBP1 has been proposed to be gradually reduced during the maturation of progenitor cells into mature neurons^82,91,93,94^. Overall, our spatial transcriptomics, augmented with selected immunocytochemistry, establishes that expression of PTBP1 and the targets it interacts with are enriched in the ependymal cells lining the SVZ ***(Figure 2N)***.

### Comparing MERFISH and single nuclear RNA sequencing

Single nuclear RNA sequencing has served as the primary tool for defining cellular identity and RNA profiles of individual cells^95^. To compare RNA profiles obtained using RNA sequencing with MERFISH methodology, we partitioned wild type, 1 year old murine brains into two halves, using the right hemisphere for MERFISH and the left hemisphere for single nuclear RNA sequencing (**Figure S2A**). Following further dissection to focus on the subventricular zone and the hippocampus, we obtained matched single nuclear RNA sequencing and MERFISH imaging for these brain areas **(Figure S2A).** Subsequently, we independently generated UMAP representations and cell-type clustering for each of the two techniques, resulting in the identification of 23 clusters through single nuclear RNA sequencing and 59 clusters through MERFISH, from a total of 54,801 pooled nuclei and 83,981 cells from a single image, respectively (***Figure S2B,C)***. Our analysis primarily encompassed the principal clusters, including neurons (dentate gyrus, GABAergic, or glutamatergic) and all anticipated glial cells (astrocytes, ependymal cells, oligodendrocytes, and microglia). After collapsing the 7 astrocytic classes, 9 oligodendrocytes, 21 neuron classes, 7 microglia, 5 ependymal, and 8 endothelial clusters identified by MERFISH into single larger cell type cluster (***Figure S2D)***, a good concordance (with up to 0.6 Pearson correlation coefficient) between the RNA signatures for each of the six clusters found between the two techniques was identified.

Additionally, we focused on a shared set of 223 genes, evaluating key parameters such as the total transcripts per cell or nucleus and the presence of genes with more than 10 transcripts per cell or nucleus in either the MERFISH or nuclear RNA sequencing data, respectively. Out of an average of 44.4 × 10^4^ reads per nucleus with single nuclear RNA sequencing, a median of only 210 total transcripts per nucleus was found for the 223 genes in our MERFISH library. In contrast, MERFISH imaging identified 1048 total transcripts per cell for the corresponding RNAs (**Figure S2E,F**). We focused on the three central clusters composed of 1) mature dentate gyrus neurons, 2) GABAergic neurons, and 3) ependymal cells and three genes of particular interest (*Prox1* for the dentate gyrus, *Gad1* for GABAergic neurons, and *Ptbp1* for ependymal cells (**Figure S2G,H**). Cross comparison of expression levels for the three genes in each of the three cell clusters revealed that MERFISH detected RNAs for both abundant RNAs (such as *Prox1* and *Gad1* in dentate gyrus and GABAergic neurons, respectively) and less abundant RNAs (e.g., *Ptbp1*), while at the read depth captured by our typical single nuclear analysis was not powered to quantify lowly expressed genes (e.g., *Ptbp1*) (**Figure S2G,H**).

To further validate the MERFISH results, we used single-molecule FISH (smFISH) to quantify the expression levels of the 223 MERFISH genes in an 8-week-old mouse. We designed the smFISH probes targeting each gene to contain two unique binding sites such that each gene could be independently quantified twice in each cell **(Figure S3A).** We also covalently linked the smFISH probes in a thin polyacrylamide gel casted on top of the sample to allow many rounds of hybridization without signal degradation **(Figure S3A)**. Across 75 rounds of hybridization and 3-color imaging we established that *1)* the smFISH results did not degrade upon the many cycles of hybridization **(Figure S3B)** and *2)* the results were quantitively consistent with those from MERFISH **(Figure S3E,F,G,H).**

### Transient ASO-mediated suppression of Ptbp1 induces activation of pro-neuronal genes

In line with our previous work^46^, one-year-old mice were administered (via intracerebroventricular (ICV) injection) with an antisense oligonucleotide (ASO) targeting the 3’ untranslated region of the mRNA encoding PTBP1. Two weeks post-treatment, brain tissue was collected and *Ptbp1* mRNA levels assessed **(Figure 3B,C)** by smFISH and MERFISH, utilizing a panel of 50 probes targeting *Ptbp1* mature transcripts. As expected, *Ptbp1* RNAs were reduced by 40%-50% across all brain regions examined. This reduction was most prominent in the hippocampus and subventricular zone **(Figure 3C,D)**, encompassing astrocytes, neurons, choroid plexus, and ependymal cells **(Figure 3E,F,G,H)**. Ptbp1 transcript levels within hippocampal and ventricular astrocytes, respectively, were reduced to 37% and 46% compared to controls **(Figure 3E,F)**, with similar decreases observed in CA1, CA3, and DG hippocampal neurons **(Figure S4)**. RNAs previously proposed to be PTBP1 targets^82,91,92^ **(Figure 2J),** including those encoding REST and RTN4, remained at comparable levels in each of these cell types **(Figure 3E,F).** In contrast, RTN4 and REST were reduced (with an average reduction of 63% and 48%, respectively) in cells of the choroid plexus and ependymal cells lining the subventricular wall that were highly expressing *Ptbp1* **(Figure 3G,H)**. Notably, *Dcx*, an established marker of immature neurons in adult neurogenesis^70,96^, was induced, albeit weakly, only within ependymal cells among the glial cell types investigated **(Figure 3H)**.

**Figure 3:**
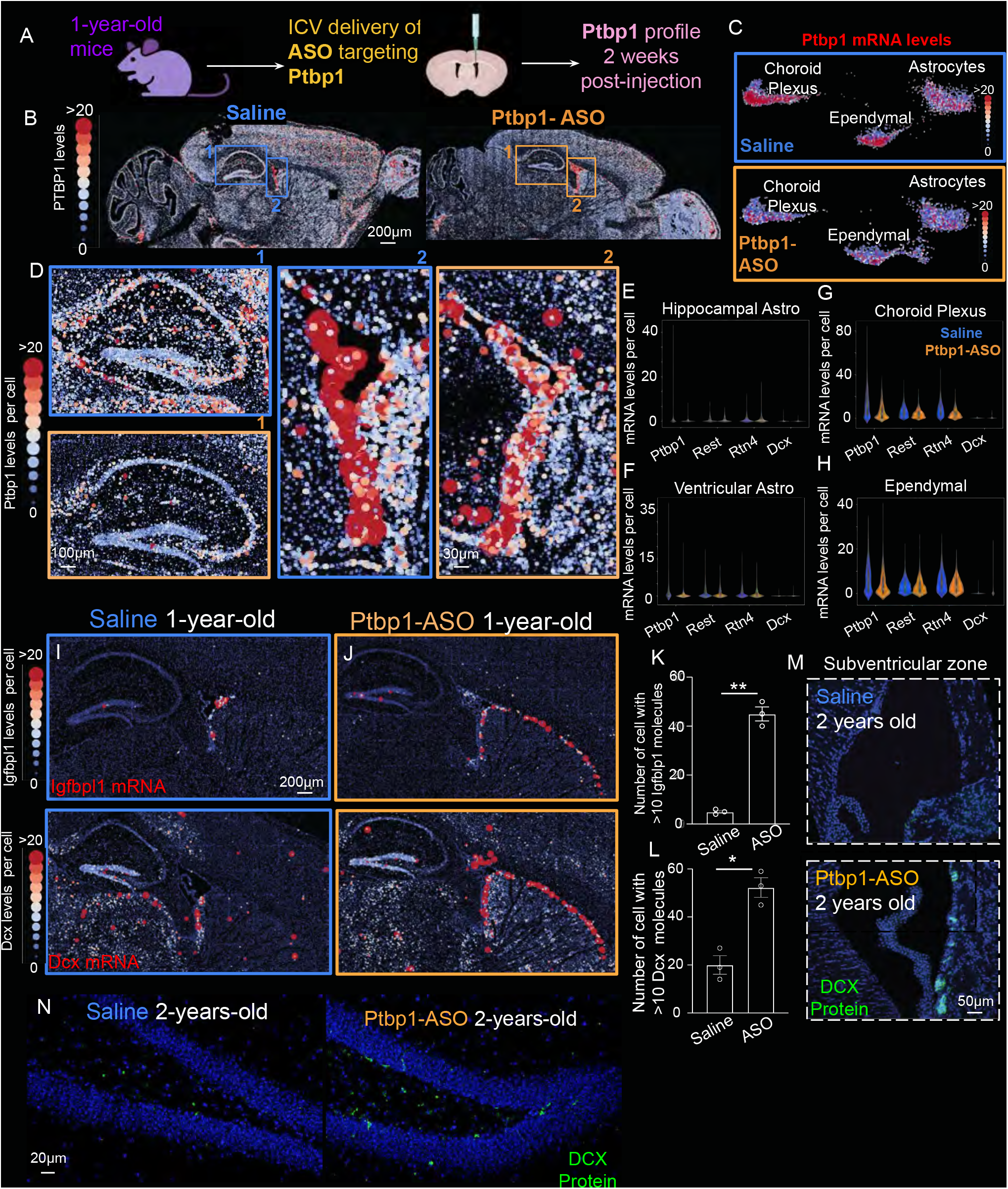
PTBP1 suppression modulates gene expression in the aged mouse brain. **(A)** Schematic of the experimental design illustrating the delivery of Ptbp1-ASOs throughout the cerebrospinal fluid (CSF) of 1-year-old mouse brain. The levels of *Ptbp1* mRNA and other genes were quantified 2 weeks post-injection using MERFISH and single-molecule FISH (smFISH). **(B,C)** Representative spatial cell maps **(B)** and partial UMAPs **(C)** showing the *Ptbp1* mRNA levels in the brain of in 1-year-old mice injected with saline (blue) or Ptbp1-ASO (orange) 2 weeks post-injection**. (D)** Insets of (B) highlighting the (1) hippocampus and (2) subventricular zone (SVZ). **(E,F,G,H)** Quantification of mRNA levels of *Ptbp1*, *Rest*, *Rtn4*, and *Dcx* in hippocampal astrocytes **(E),** ventricular astrocytes **(F),** choroid plexus **(G),** and ependymal cells **(H)** from 1 year old saline or Ptbp1-ASO treated mice**. (I,J,K,L)** Representative spatial maps quantifying single-cell expression levels (I, J) and corresponding bar graphs quantifying the number of *Igfbpl1*-expressing □=≥20 counts, (top)] and *Dcx*-expressing □=≥20 counts, (bottom)] of 1-year-old sagittal sections in saline or Ptbp1-ASO, 2 weeks post-injection (n=3, t.test **p=0.0043, t.test *p=0.03). **(M,N)** Representative images of immunofluorescence for DCX (green) in 2-year-old mouse brain in the subventricular zone **(M)** and dentate gyrus **(N)** 2 weeks post-injections of saline control (blue) or PTBP1-ASOs (orange). Blue: DAPI.

We further observed higher levels of RNAs encoding DCX and IGFBPL1 in additional cell types along the rostral migratory track connected to the SVZ and within a few cells in the SGZ of the dentate gyrus. This effect was particularly pronounced for the SVZ, with approximately 60 cells positive on average for IGFBPL1 RNAs (≥20 transcripts per cell) in the PTBP1-ASO injected group compared to 4 positive cells on average in control animals injected with saline **(Figure 3I,J,K,L)**. Complementing these findings, traditional immunofluorescence staining for DCX revealed an eightfold increase in the number of cells expressing DCX along the SVZ and a threefold increase in the dentate gyrus (DG) in even older, two-year-old animals, two weeks post-ASO delivery **(Figure 3M,N).** Collectively, our data suggest that PTBP1 suppression modulates the expression of genes involved in the neurogenesis process within these two areas of the brain.

### Adult neurogenesis in the dentate gyrus and SVZ of young mice

To better understand the different cell stages and activated genetic program in this induced process and compare it with the adult neurogenesis process in young animals, we first constructed a more detailed atlas of endogenous adult neurogenesis within young animals. We focused on cells within a sagittal section of the hippocampus of a 4-week-old mouse **(Figure 4A).** Re-clustering of 4,164 hippocampal cells (based on their MERFISH-derived single-cell transcriptomes) identified 45 cell types covering neuronal and glial populations specific to subregions of the hippocampus **(Figure 4B)**, including resident astrocytes of the CA1, CA2, or CA3 regions **(Figure 4B)** as well as small populations of cells such as the inhibitory neurons of the Hillus. The dentate gyrus was found to contain cell layers comprised of five cell groups organized along a basal-apical axis **(Figure 4C).** Mature dentate gyrus neurons, forming the most apical cluster, expressed typical markers such as *Prox1* and axonal guidance molecules including *C1ql3*.

**Figure 4:**
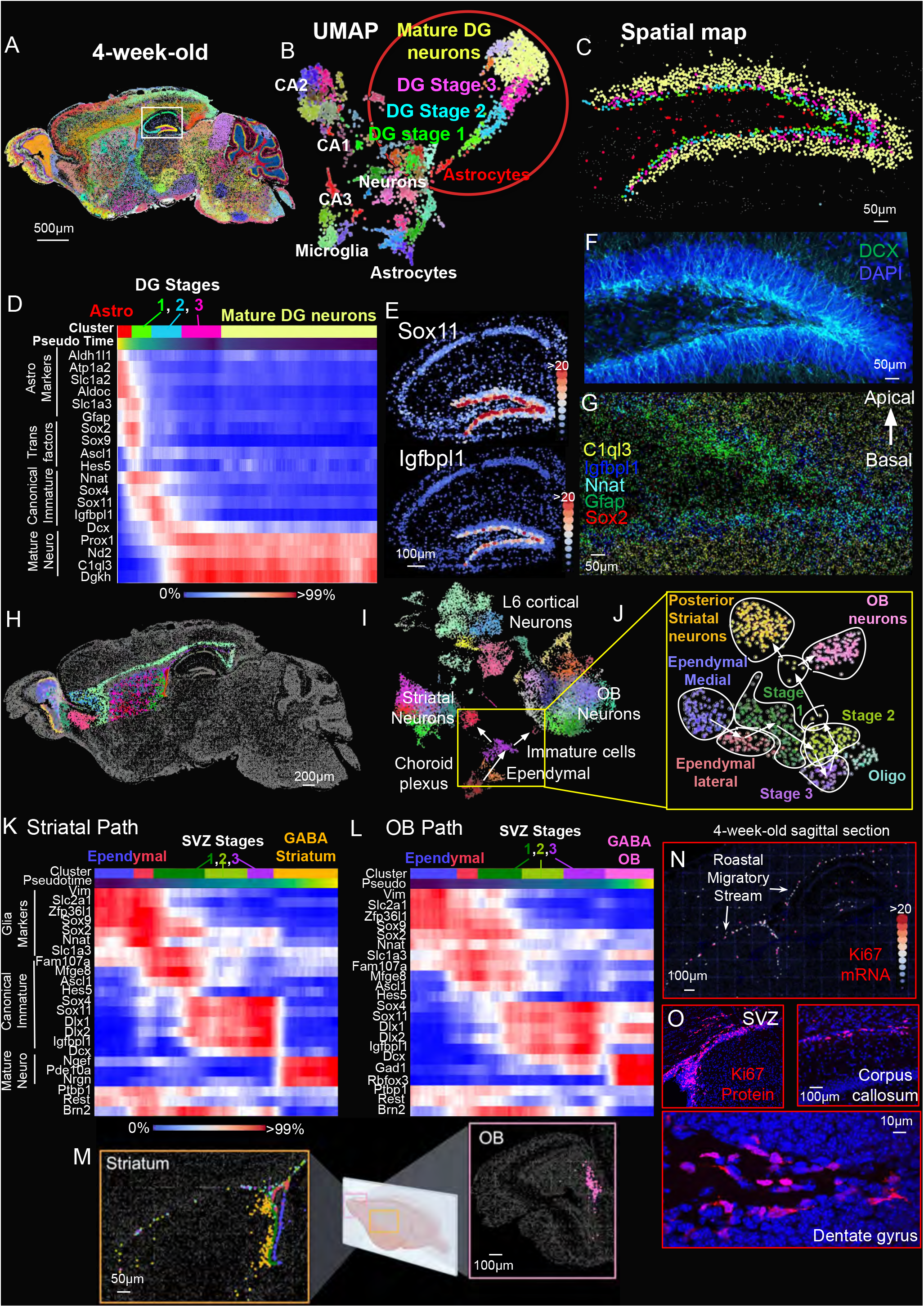
A detailed spatial and molecular atlas of adult Neurogenesis in the dentate gyrus and subventricular zone. **(A)** Representative spatial map of a 4-week-old mouse sagittal section in which cell-types identified by MERFISH are highlighted with different colors. **(B)** UMAP of the hippocampal sub-area (white square in (A))**. (C)** Spatial map of the cell clusters identified the dentate gyrus area from (B) with matching colors. **(D)** Heatmap showing the mRNA expression levels of the representative genes in each cell-type cluster marked in (B) [Astrocytes (red), DG Stage 1 (green), DG Stage 2 (blue), DG Stage 3 (pink) and Mature DG neurons (yellow)], and their dynamics along pseudo time transcriptional space. **(E)** Representative spatial maps quantifying the expression levels per cell of *Sox11* (top) and *Igfbl1* (bottom) in the hippocampus brain region of a 4 week-old brain. **(F)** Representative image of immunofluorescence (IF) of DCX (green) in a 4-week-old dentate gyrus. DAPI (blue). **(G)** Representative image of individual mRNAs detected through MERFISH of *Sox2* (red), *Gfap* (green), *Nnat* (white), *Igfbpl1* (blue), and *C1ql3* (yellow) along dentate gyrus horns **(H)** Representative spatial map of the cell clusters identified by MERFISH in the ventricular zone, the striatum and olfactory bulb of a 4-week-old mouse brain. **(I)** UMAP of the clusters in (H). **(J)** UMAP for the cells connected along the neurogenesis pathways in the subventricular zone (SVZ). **(K,L)** Heatmap showing the mRNA expression levels of the representative genes in each cluster of the SVZ along the striatal **(K)** and olfactory bulb **(L)** neurogenesis paths [Ependymal (blue, red), SVZ Stage 1 (green), SVZ Stage 2 (light green), SVZ Stage 3 (purple) and GABA neurons (yellow), GABA OB (pink)]. The cells are ordered along the pseudo time defined in transcriptional space. **(M)** Representative spatial maps of the cell clusters in (K) and (L) **(N)** Representative spatial map quantifying the single-cell expression levels of *Mki67* in a 4-weeks sagittal section. **(O)** Representative images of IF for KI67 (red) in the DG and SVZ of a 4-week-old mouse.

Starting from mature dentate gyrus neurons, we computationally defined a pseudo-time axis by progressively connecting the mature neurons to their closest neural progenitors in transcriptional space using a single-cell analysis package called PAGA (see Methods). Along this continuous pseudo-time axis, we captured the following neurogenic clusters in order of maturation: radial-glia like cells (DG Stage 1), early immature neurons-like cells (DG Stage 2), late immature basal neurons (DG Stage 3), and mature apical DG neurons **(Figure 4B,C),** consistent with prior single-cell sequencing reports in which similar cell stages and maker genes for each stage were observed along the adult neurogenesis process in the dentate gyrus^70^ **(Figure S5A-C)**. The high detection efficiency of mRNA molecules with the MERFISH methodology further allowed the determination of the approximate temporal progression of gene expression during this neuronal maturation process **(Figure 4D)**. RNAs encoding transcription factors *Sox2*, *Sox9*, *Ascl1* and *Hes5* were found to be expressed by radial glia-like cells in the most basal layer of the dentate gyrus **(Figure 4D)**. Three of these (*Sox9*, *Ascl1* and *Hes5*) were silenced prior to DG Stage 2 cells, while *Sox2* expression continued within the immature neuronal states, consistent with prior studies **(Figure 4D)**^71,82,91,93,94^. Within the later immature states, there was transient cascade of genes induced including *Sox4*, *Sox11* and then *Dcx* **(Figure 4D).** Finally, as the new neurons matured, genes encoding axonal guidance molecules and neuropeptides specific for their function (e.g., *Scla7* and *C1ql*) were expressed ***(Figure 4D*).** This temporal progression was also mirrored in physical space, in which the same transcription factors and axonal guidance molecules/neuropeptides progress along the basal-apical direction of the dentate gyrus **(Figure 4E,F,G).**

We applied the same approach for the SVZ, starting with re-clustering more finely the 17,699 cells around the lateral ventricle, the rostral migratory path, the striatum, and the olfactory bulb ***(Figure 4H,I,J).*** We focused on the cell types with immature neuronal character (e.g., cells expressing *Sox11* and *Dcx*). We determined which cell types they were most closely connected to among glia as a means to identify their origin, and which cell types they became most closely related to among the neuronal population, suggestive of their final destinations and identities. Upon applying PAGA analysis as we had done for the dentate gyrus, these immature cells were identified to be most closely connected to a subgroup of ependymal cells, which we named SVZ Stage 1 cells **(Figure 4K,L)**. These cells were similar to the other ependymal cells expressing vimentin and the glutamate transporter Glast, but also had radial-glia like character (expressing *Ascl1* and *Hes5*) ***(Figure 4K,L)***. Immature cells with neuronal character were connected along two paths, either mature neurons in the posterior regions of the striatum (near the lateral ventricle) or mature granule neurons in the olfactory bulb **(Figure 4K,L,M).** These branches of connectivity in transcriptional space are consistent with previously reported tracing experiments performed in young animals in which ependymal cells were observed to undergo cell proliferation, initiating expression of immature neuronal markers, and incorporation into the olfactory bulb and, to a lesser extent, into the striatum^97^.

Among the genes modulated along this neurogenesis conversion process, *Mki67* and *Top2a* expression was initiated in SVZ Stage 1 immature cells ***(Figure 4N,O),*** suggesting that these cells undergo cell division, consistent with prior studies in which DNA synthesis (marked with BrdU) was used to identify dividing cells^98^. Comparison of the cell types we identified with the data from a prior single nuclear sequencing effort within the SVZ^71^ yielded a good correspondence (e.g., R = 0.62 for mature DG neurons) between gene expression observed in their data and ours **(Figure S5D,E,F)**.

### Induction of immature proliferating cells following transient suppression of Ptbp1 in the neurogenic niches of the aged mouse brain

We next tested the consequence of antisense oligonucleotide (ASO) mediated, transient PTBP1 suppression within the dentate gyrus and subventricular zone of 1-year-old mice utilizing the comprehensive atlases of neurogenesis in young (4 week old) mouse brain as initial reference points **(Figure 4)**. Compared with young (4 weeks or 8 weeks) brains, neurogenesis was nearly eliminated in 1 year old brains, consistent with our earlier evidence **(Figure 1J)** and previous reports^99,100^. Specifically, within the dentate gyrus of a wildtype 1 year old mouse, while radial glia-like cells (DG stage 1) persisted at half the frequency of young mice, neuronal progenitor cells and immature neurons (DG stages 2 and 3) were almost eliminated (with a 30 fold drop in their number in the 1 year old untreated brain compared to that in young mice) **(Figure S6A).** After two weeks of PTBP1 suppression, the number of DG stage 1 cells remained largely unchanged compared to saline injected or non-injected aged animals. DG stage 2 cells displayed a modest increase in numbers, while DG stage 3 cells displayed a 2 fold increase ***(Figure S6B,C).*** While the total number of these transient DG stage 3 immature cells, captured at a single timepoint, remains small, the fold increase upon PTBP1 suppression is roughly consistent with the number of new neurons labeled with the GFAP:CRE genetic strategy in 1 year old treated animals in our prior study in which we observed an increase of labeled new neurons by ∼4 fold over the course of PTBP1 suppression for 2 months^46^. Performance in a hippocampal dependent test^101^ (the Barnes maze) significantly improved in PTBP1 injected groups of 1 or 2 year old mice, compared with age and sex matched control groups **(Figure S6D**), consistent with our initial report^46^.

Similarly, in the aged (1 year old) SVZ, there was a sharp decrease in subependymal-like cell types (SVZ stage 1, SVZ stage 2, and SVZ stage 3) compared with 4 weeks old control mice **(Figures 1J, 5A)**. Despite a mostly unchanged number of SVZ stage 1 cells, 2 weeks post PTBP1 suppression there was a 2.3 fold increase in the number of SVZ stage 2 cells and an 8-fold increase in the SVZ stage 3 cells compared with non-injected or saline injected controls **(Figure 5A,B)**. Comparison (PCA analysis) of gene expression profiles within these SVZ stage 1 cell populations of treated and control groups identified that ASO-mediated reduction in PTBP1 both induced canonical genes of immature neurons (including *Dcx*, *Sox11* and *Ascl1*) and suppressed genes typically considered markers of mature glia (including *Vim*, *Gja1*, *Aqp4*) **(Figure 5C,D)**. Additionally, genes that are markers of cell proliferation^102,103^(including *Mki67*, *Ccnd1*, *Ccnd2*, and *Mcm5*) were increased **(Figure 5C,D,E)** within this cell population. The transcriptomes of SVZ stage 3 ASO induced cells were very similar to the corresponding SVZ stage 3 neurons of young animals, with no noticeable separation in PCA space **(Figure 5F,G)**.

**Figure 5:**
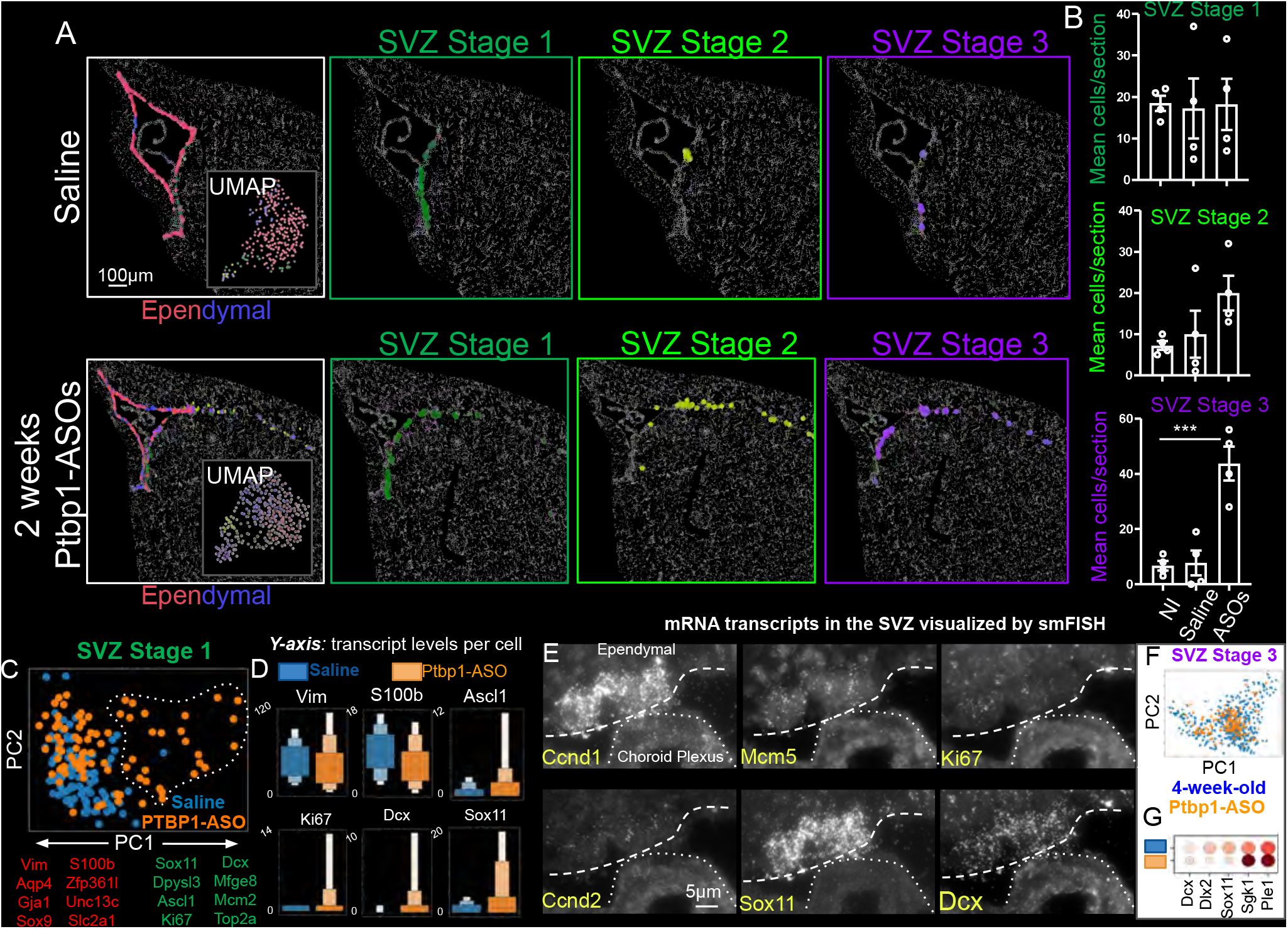
Ptbp1-ASOs Intra-cerebro-ventricular delivery induced activation of quiescent subependymal cells. **(A)** Representative spatial map and UMAP of 1 year-old mice 2 weeks post injection with saline (upper) and Ptbp1-ASOs (lower), highlighting the clusters: Medial Ependymal (blue), Lateral Ependymal (red), SVZ-Stage 1 (green), SVZ-Stage 2 (light green), SVZ-Stage 3 (purple). **(B)** Quantification of the number of cells identified in ‘SVZ Stage 1’, ‘SVZ Stage 2’ and ‘SVZ stage 3’ cell types in non-injected 1 year-old mice and 2 weeks post saline or ASO injection, respectively (n□=□4 animals per each condition; one-way ANOVA, ***p□=0.0004□). **(C)** Principal component analysis (PCA) of SVZ-Stage 1 cells from 1 year-old saline and PTBP1-ASO injected mice. The top differentially expressed genes along the PC1 axis are highlighted. **(D)** Quantification of the mRNA levels of *Vim, S100b, Ascl1, Mki67, Dcx* and *Sox11* genes in SVZ Stage 1 cell clusters of 1 year-old mice, 2 weeks post saline or ASO injection. **(E)** Representative single molecule FISH images in the SVZ of a 1 year-old mouse 2 weeks post ASO injection for *Ccnd1, Mcm5, Mki67, Ccnd2, Sox11* and *Dcx*. **(F)** PCA plot of SVZ-Stage 3 cells from 4 week-old (blue) and PTBP1-ASO injected 1 year-old (orange) mice. **(G)** Dot plot of the 5 highest expressed genes in SVZ stage 3 in 4-week-old and PTBP1-ASO injected 1 year-old mice.

### Ptbp1 suppression-induced neurogenesis reactivates DNA synthesis in dormant precursor glia

Possible cell cycle re-entry and subsequent DNA replication during re-activation of neurogenesis were evaluated by daily injection for two weeks of a thymidine mimetic (5-Ethynyl-2’-deoxyuridine (EdU)^104^) after injections in one year old saline or PTBP1-ASO injected mice or a 4-week-old wild type mouse in which active neurogenesis was still ongoing **(Figure 6A)**. After an additional two-weeks without EdU administration to permit neuronal differentiation, in all conditions the highest percentage of EdU labeled cells was found in the olfactory bulb, consistent with active and continuing neurogenesis expected for an active niche^105^**(Figure 6B,C,D)**. Counting nuclei across sagittal sections revealed that 3.8% EdU+ cells (5598 EdU+ cells out of 147,402 cells examined) in the two well established neurogenic niches (the DG and SVZ) of 4-week-old brain were EdU+, consistent with the expected DNA replication in canonical neurogenesis^81^. The number of EdU labeled nuclei was reduced almost 10 fold (to 0.4%) in 1-year-old saline-injected animals (562 EdU+ cells out of 145,607) **(Figure 6B,C,D)**. ASO-mediated transient suppression of PTBP1 in 1-year-old mice reinduced strong EdU labeling across entire individual nuclei, consistent with DNA replication not simple DNA repair in the previously dormant DG and SVZ neurogenic niches and with the frequency of such EdU+ nuclei rising to 70% of the level found in young brains (3834 EdU+ cells out of 143,582 or 2.7%) **(Figure 6B,C,D)**. In striking contrast, almost no (<3) cells in either niche were EdU labeled in 1 year old saline injected mice.

**Figure 6:**
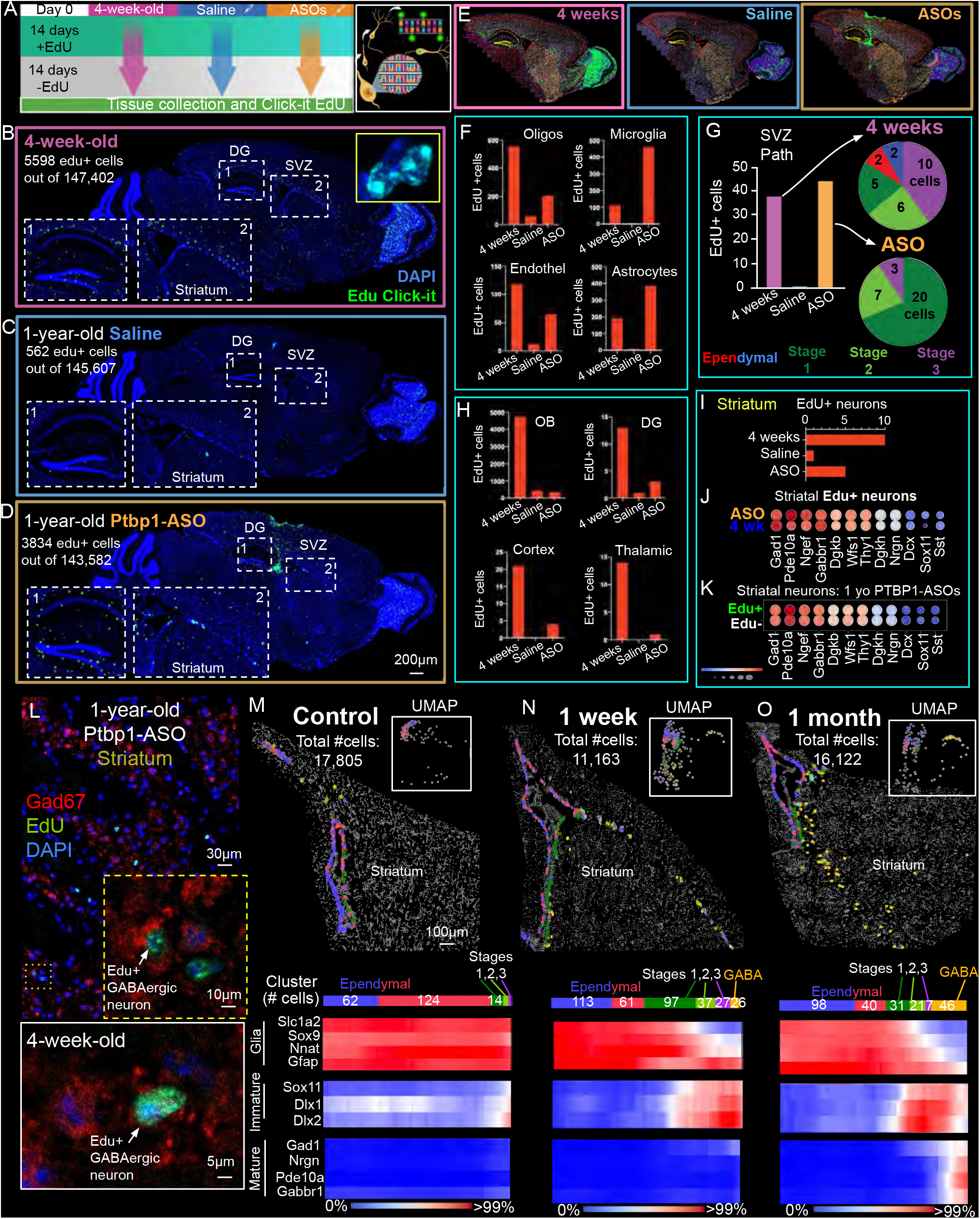
Tracing newly generated neurons by coupling MERFISH spatial transcriptomics with Edu labeling. **(A)** Schematic of the DNA-labeling strategy used to target proliferating cells upon EdU incorporation throughout the brain. EdU was administered by daily intra-peritoneal injections for 2 weeks to a 4-week old mouse and 1-year-old mice saline or Ptbp1-ASO injected. Brains across all conditions were collected 1 month from the start of the treatment and EdU was fluorescently tagged using Click chemistry. **(B,C,D)** Representative images and corresponding insets of 4 week **(B)** 1 year old saline **(C)** and 1 year old PTBP1-ASO **(D)** EdU-labeled brains. EdU (green), DAPI (blue). **(E)** MERFISH spatial maps of the cell types identified in 4-week-old (left), 1-year-old saline (middle) and Ptbp1-ASO (right) mice. EdU positive cells were highlighted in green. **(F)** Quantification of EdU-positive cells in each of the three experimental groups within major non-neuronal cell types including Oligodendrocytes, Microglia, Endothelial cells and Astrocytes. **(G)** Quantification of the number of EdU-positive cells in each of the three experimental groups across the cell-types part of the SVZ neurogenesis path including ependymal cells and SVZ stages 1,2 and 3. **(H)** Quantification of the number EdU-positive cells in each of the three experimental groups in neurons including olfactory bulb (OB), dentate gyrus (DG), cortex and thalamic brain regions. **(I)** Quantification of the number of EdU-positive cells in each of the three experimental groups in striatal neurons. **(J)** Gene expression dot plots comparing EdU-positive striatal neurons of 4 week-old mice with EdU-positive striatal neurons of 1-year-old Ptbp1-ASO injected mice **(K)** Gene expression dot plots comparing EdU-positive and EdU-negative striatal neurons in 1-year-old Ptbp1-ASO injected mice. **(L)** Top: Representative immunofluorescence image of GAD67 (marker of GABAergic neurons) (red) together with EdU signal (green) in the striatum of 1-year-old PTBP1-ASOs injected mouse. DAPI in blue. Bottom: Same as top for the striatum of a wildtype 4-week-old mice. **(M,N,O)** Spatial maps, UMAPs and gene expression profiles of cells connected across the SVZ neurogenesis path in 1 year-old mice injected with saline **(M),** or **(N)** 1 week- and **(O)** 1 month post PTBP1-ASO delivery.

The identity(ies) of the EdU+ cells appearing after PTBP1 suppression were determined by combining EdU labeling and imaging with MERFISH in the same single cells. Globally, across all the three conditions tested and all brain regions profiled (including hippocampus, SVZ, thalamus, midbrain, cortex and olfactory bulb (OB)) **(Figure 6E)**, the four major cell types undergoing EdU labeling were found to be microglia, astrocytes, oligodendrocytes, and endothelial cells **(Figure 6F)**. The number of cycling cells in the 1 year old animals fell to ∼1% the level seen in 4 week old brains, while single injection of PTBP1-ASO in 1 year old mice induced proliferation of all glial and endothelial types **(Figure 6F).** Specifically, in single sections within the SVZ of 4 week old mice **(Figure 6G)**, EdU+ cells included a small number of ependymal cells and ∼20 immature cells that were identified with MERFISH to be at SVZ stages 1, 2 or 3. In 1-year-old saline-injected mice, there was almost a complete absence of EdU+ cells (only 1 per section in the SVZ). Remarkedly, PTBP1 ASO suppression resulted in restoration of a similar number of labelled immature SVZ stage 1, 2, or 3 neurons as was found in 4 week old mice, consistent with reactivation of the SVZ neurogenic niche **(Figure 6G).**

Among neuronal types, the OB exhibited the highest number of labelled neurons in 4-week-old mice, which decreased by almost 90% in 1 year old mice (saline or PTBP1-ASO injected). PTBP1 ASO suppression did not produce a significant increase in EdU+ neurons in the OB of 1-year-old mice **(Figure 6H)**. In contrast, compared with the 1 year old saline injected mice, PTBP1 suppression induced increased numbers of EdU+ neurons across multiple brain areas (including the cortex, DG, midbrain, and thalamus) **(Figure 6H)**, with the highest fold increase in the striatum, with ASO injection increasing the number of EdU+ neurons to about half of that in 4 week old mice **(Figure 6I)**. Transcriptional profiling of EdU+ striatal neurons within a 4 week old mouse and a 1 year old mouse injected with PTBP1-ASO revealed that these neuronal populations were highly similar, with both expressing the expected striatal GABAergic neuronal markers **(Figure 6J)**. Similarly, 1 month after PTBP1 ASO injection, EdU+ (newly made) and EdU-(preexisting) striatal neurons in 1 year old mice injected with PTBP1-ASO were highly similar transcriptionally, with high levels of *Gad1*, *Pde10a*, and *Ngef*, while markers of immature neurons (including *Dcx* and *Sox11*) were low **(Figure 6K)**. Contemporaneous antibody staining and imaging for Gad67 and EdU confirmed the presence of newly generated inhibitory neurons in the striatum **(Figure 6L)**.

To better characterize the process underlying induced neurogenesis, the ***1)*** number of cells for each cell type and ***2)*** the gene expression along the induced neurogenesis trajectory were quantified at two different time points (1 week and 1 month) post PTBP1-ASO injection **(Figure 6M,N,O).** A transient increase in SVZ stage 1 cells was detected within 1-week of PTBP1 suppression, but returning to baseline within 1 month after ASO injection. The number of SVZ Stage 2 and SVZ stage 3 cells increased at the 1-week time point, then declined in abundance by the end of 1 month. Tracing individual genes across pseudo-timepoint trajectories (using the PAGA algorithm) revealed the same wave of neurogenic gene induction in PTBP1-ASO injected aged mice as was found in natural neurogenesis in the 4-week old brain. For instance, in both young and aged mouse brains, *Dlx1* and *Dlx2* appeared in SVZ Stage 2 and SVZ Stage 3 cells, but *Nrgn*, *Gad1* and *Pde10a* induction only appeared in maturing GABAergic neurons. Altogether, PTBP1 suppression in the aged mouse brain induces a canonical neurogenesis-like process initiating within an ependymal-like cell subpopulation (SVZ stage 1), involving cell division and proliferation and resulting in GABAergic neurons in the striatum with transcriptomes similar to preexisting GABAergic neurons.

### Ependymal cells highly expressing PTBP1 in the SVZ of aged human and non-human primates

To test whether a similar class of PTBP1-expressing ependymal cells lining the ventricle exists in aging human brain, we implemented MERFISH on human postmortem brain tissue containing part of the lateral ventricle and striatum from an aged (68 yr old) individual **(Figure 7A)**. We generated a 1130 human gene list including 357 genes for cell typing selected based on the Allen’s Institute’s single nuclear RNA sequencing (snRNAseq) dataset for human brain, genes covering multiple transcription factor classes^78^ (e.g., *PAX6*, *SOX2*, etc.), genes expressed within cell types within neurogenic niches^79,80^ (e.g., *AQP4* and *GFAP* marking astrocytes; *DCX* and *NEUROD1* marking immature neurons; *FOXJ1* marking ependymal cells), and genes within or related to the proposed PTBP1 pathway^82–84^ (e.g., *PTBP1*, *PTBP2*, *REST*, *CLASP1*) ***(Figure S8)***. Based on the RNA contents **(Figure 7B)** of each imaged cell, unsupervised clustering (using the Leiden algorithm^87^) identified 41 cell type clusters from the 18,364 cell transcriptomes within a single image. We first annotated each of these clusters into the 9 major cell-type classes (these included glutamatergic and GABAergic neurons, two classes each of astrocytes and oligodendrocytes, microglia, ependymal and endothelial cells) based on the expression of the expected marker genes **(Figure 7C,D)**. The cell types identified had a distinct spatial organization within the section (**Figure 7C,D).** The different subtypes of neurons displayed the expected localization within the striatum, while ependymal cells lined the ventricle walls and oligodendrocytes appeared mostly in the white matter. A Pearson correlation heatmap comparison of our identified clusters in the human and murine brains **(Figure 7E)** identified with MERFISH indicated a high degree of similarity (e.g., a Pearson correlation of 0.61 comparing neurons).

**Figure 7:**
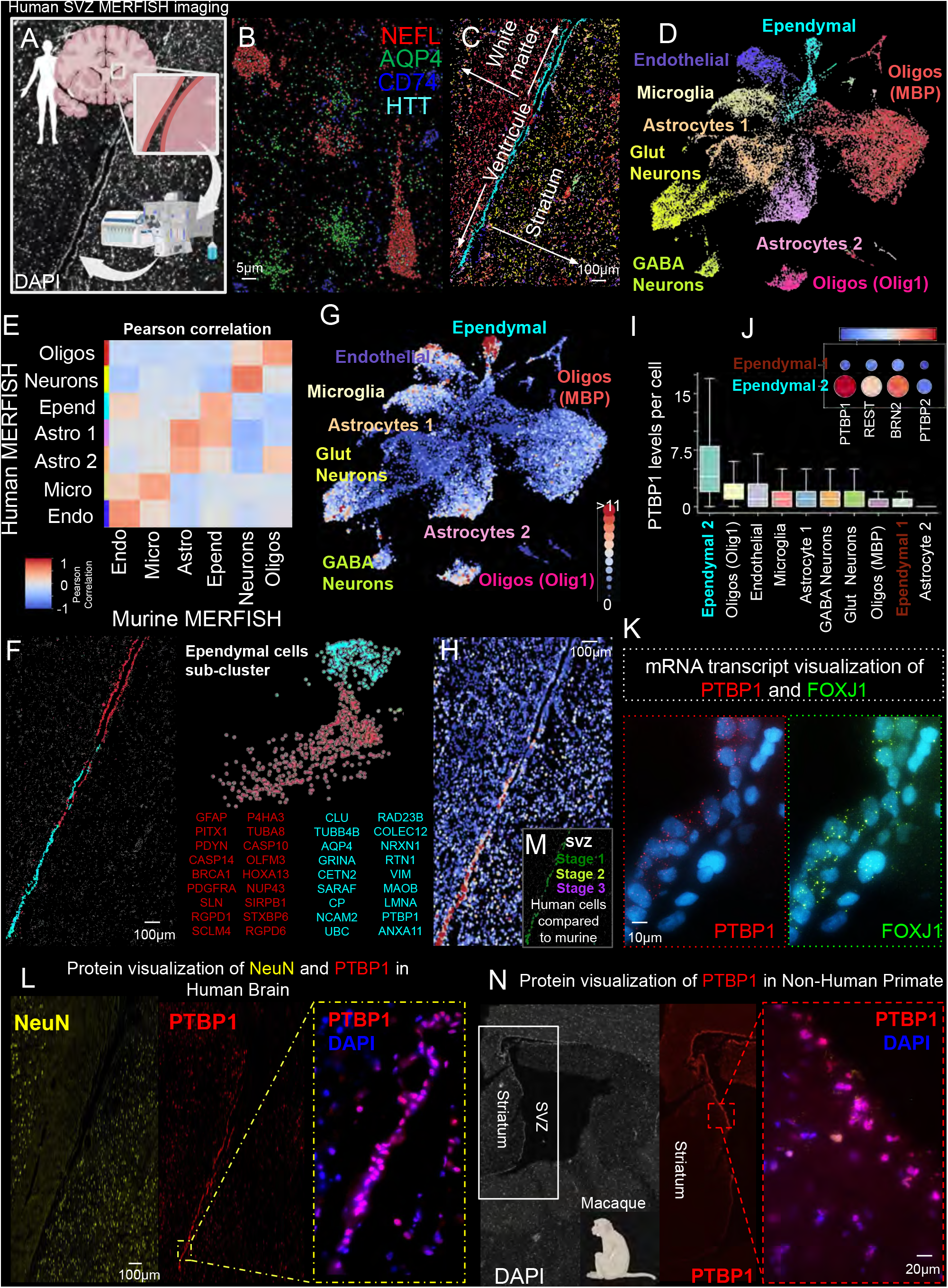
PTBP1 and its proposed regulatory loops are expressed in the SVZ of human and non-human primates. **(A)** Schematic of the human brain region from which the SVZ was dissected and imaged via MERFISH for ∼1200 genes. **(B)** Image of MERFISH-decoded mRNAs for *NEFL* (in red, neuronal marker), *AQP4* (in green, astrocytic marker), *CD74* (in blue, microglial marker), *HTT* (in cyan, neuronal gene). **(C**, **D)** Spatial map and UMAP of the cell types identified in the SVZ human tissue section. **(E)** A Pearson correlation heatmap comparing the major cell type transcriptional profiles quantified by MERFISH in the human and mouse SVZ. **(F)** Spatial map (left) and UMAP (right) of ependymal cell sub-clusters identified in the human SVZ. The genes with the highest differential gene expression in each subcluster are highlighted. **(G)** *PTBP1* levels per cell depicted in the UMAP of the human SVZ. **(H,I)** The spatial map **(H)** and box plot quantification **(I)** of *PTBP1* levels per cell in the human SVZ. **(J)** Dot plot of *PTBP1* and PTBP1-controlled genes (*PTBP1*, *REST*, *BRN2*, *PTBP2*) expression in Ependymal cells 1 and Ependymal cells 2 clusters. **(K)** Representative single molecule FISH images of *PTBP1* (red) and *FOXJ1* (green, ependymal cell marker) in the human SVZ. DAPI in blue. **(L)** Representative immunofluorescence (IF) of NeuN (yellow, neuronal marker) and PTBP1 (red) in the human SVZ. DAPI is depicted in blue. **(M)** Spatial map of SVZ Stage 1,2 and 3 human cell types as inferred upon comparing with the murine brains. **(N)** Representative images of IF of PTBP1 (red) in the non-human primate SVZ. DAPI in blue.

Among the ependymal cells (cyan, in **Figure 7F**) identified within the MERFISH images of human brain, we noticed two physically and transcriptionally distinct sub-populations. After sub-clustering these cells, one ependymal cluster lined the top of both sides of the SVZ (**Figure 7F**) and was characterized by low *PTBP1* expression, with high expression of *GFAP*, *PITX1*, *PDYN*, *CASP14*, *BRCA1*, *PDGFRA*, *SLN*, *RGPD1*, *SCLM4*, *P4HA3*, *TUBA8*, *CASP10*, *OLFM3*, *HOXA13*, *NUP43*, *SIRPB1*, *STXBP6*, and *RGPD6* genes. The second ependymal cluster lined the bottom of the SVZ (**Figure 7F**) and was characterized by high expression of *PTBP1*, as well as *CLU*, *TUBB4B*, *AQP4*, *GRINA*, *CETN2*, *SARAF*, *CP*, *NCAM2*, *UBC*, *AD23B*, *COLEC12*, *NRXN1*, *RTN1*, *VIM*, *MAOB*, and *LMNA* **(Figure 7F)**. Indeed, in this second ependymal cluster *PTBP1* was the 15^th^ most abundant RNA of the RNAs quantified, accumulating to a level that was the highest of the 41 human cell type clusters identified **(Figure 7F)**.

Clustering identified two astrocytic classes in the region imaged, one located in the white matter with a low level of *PTBP1* encoding RNAs and the other in the striatum with a nearly undetectable level of PTBP1. Endothelial cells, oligodendrocytes, and glutamatergic neurons identified by MERFISH also accumulated low or very low levels of *PTBP1* RNAs (**Figure 7G,I**). For each cell type, genes expected to depend on *PTBP1* (including *REST* and *POU3F2*) were strongly correlated with *PTBP1* in ependymal cells, but *PTBP2* level was negatively correlated (**Figure 7J**). Comparison of the human cells with the murine cells highlight the presence of quiescent SVZ stage 1 cells in the human SVZ (**Figure 7M**) and the complete absence of SVZ stage 2 and 3 cells. Use of smFISH ***(Figure 7K)*** and measurement of PTBP1 protein level ***(Figure 7L)*** confirmed that RNAs encoding *PTBP1* or *FOXJ1* were highest in ependymal cells located within the lowest region of the ventricle, consistent with a central role of PTBP1 in setting or maintaining cell identity in these cells. PTBP1 protein accumulation was also confirmed in the lower part of the non-human primate SVZ as measured by immunofluorescence ***(Figure 7N)***.

## Discussion

Use of spatial transcriptomics in aging mouse and human brain, augmented with selected immunocytochemistry (including in non-human primates), has identified a class of PTBP1-expressing ependymal cells lining the human and murine ventricles. Additionally, using transient, partial suppression of *Ptbp1* (via a single-dose of ASO delivery into CSF), we have demonstrated successful re-activation of neurogenesis within two otherwise quiescent neurogenic niches of the aged (≥1 year old) mouse brain: the SGZ and the SVZ. High *Ptbp1*-expressing ependymal cells resident in the SVZ convert from a dormant neurogenic transcriptome signature into one expected for activation of the canonical adult neurogenesis pathway. Additionally, a subset of *Ptbp1*-expressing radial glia-like cells in the subgranular zone of the dentate gyrus convert into immature neurons, consistent with ultimately becoming a mature neuron as initially proposed^46^. Use of MERFISH imaging, coupled with EdU labeling to permanently mark cells undergoing DNA replication, further demonstrated that the initial events in the SVZ include 1) DNA synthesis (following presumed cell cycle re-entry and proliferation for one or more cycles) and 2) acquisition of early neuronal character including transient expression of markers of canonical neurogenesis genes (e.g., *Sox11*, *Igfbpl1*, and *Dcx*) ending days/weeks later with their maturation into striatal GABAergic neurons.

While prior studies^51,58^ reported that PTBP1 suppression can lead to generation of new neurons, all but one of these reports^46^ was limited by ***1)*** exclusive use of genetic labeling strategies that may have inadvertently labeled populations of endogenous neurons (as subsequently carefully documented by Zhang and colleagues^52^), ***2)*** the absence of evidence at intermediate stages of the proposed conversion process, and ***3)*** use only of young (8 week old) mice in which neurogenesis is ongoing. We have overcome these limitations by examination of aged mice (≥1 year old) in which endogenous neurogenesis has been almost completely silenced (**Figures 1,4**)^106^ and use of spatial transcriptomics has enabled both identification of the dormant glial precursor cells that can transition into new neurons in the aging nervous system and capture of the neuronal proliferation process through a stream of transcriptionally related cells in the brain that gradually transition both in gene expression and physical space from *Ptbp1*-highly expressing radial glia-like progenitor cells in the dentate gyrus (which sharply decrease during aging^106^) or ependymal cells lining the lower SVZ, respectively, into mature neurons some of which migrate into the striatum (**Figure S4)**.

Our evidence is in contrast with prior claims that *Ptbp1* or *PTBP1* suppression induces mature astrocyte-to-neuron conversion^51^. Three lines of evidence suggest most new neurons induced stem from a subtype of subependymal cells (defined here as Stage 1) lining the ventricle wall after *PTBP1/Ptbp1* is suppressed: **1)** PTBP1 and its proposed targets are highly expressed in these cells in untreated aged mice; **2)** upon treatment with a *Ptbp1* targeting ASO, PTBP1 expression and that of its target genes is altered in these cells (**Figure 3**); and **3)** upon ASO administration, the subependymal cells connect in the transcriptional space to striatal GABAergic neurons (**Figure 5**). Mature astrocytes seem an unlikely cell type to be the cell of origin of *PTBP1/Ptbp1* depletion-induced neurogenesis as we have documented they initially express only low PTBP1 levels (quantified here via MERFISH and smFISH (**Figure 2**)) and treatment with a PTBP1-targeting ASO does not yield further detectable decrease in expression of other genes in the *Ptbp1* loop (**Figure 3**). Consistent with this view, recent lineage tracing experiments that labeled mature astrocytes via Alh1l1 promoted Cre concluded that indeed mature astrocytes are not likely convert to neurons upon *Ptbp1* depletion^56^.

The intermediate stages within the conversion process have been identified from the starting glial population to the sub-neuronal types generated. In contrast to single nuclear sequencing whose shallow read depth does not provide a complete transcriptome for each individual nucleus, we gained not only spatial insights but also achieved higher per cell transcriptional resolution, enabling more precise quantification of low-expressing genes crucial to characterize both the endogenous and induced neurogenic process(es) (**Supplementary Figure 3)**.

Taken together, our findings demonstrate that activation of quiescent neurogenic niches of aging brain can be achieved with a therapeutically feasible approach of ASO-mediated transient reduction in *Ptbp1*. Moreover, we identified in the aging human brain a similar class of PTBP1-expressing ependymal cells (SVZ Stage 1) lining the ventricle (**Figure 6**), suggesting the promise of re-activation of neurogenesis in humans. The reduction of PTBP1 is a particularly attractive therapeutic intervention, since ASO injection is a current standard of care for the fatal childhood disease spinal muscular atrophy^107^ and an inherited form of ALS from mutation in superoxide dismutase^108^, and is in at least five ongoing clinical trials in amyotrophic lateral sclerosis (ALS), Parkinson’s, and Alzheimer’s diseases^109^. That said, the ASOs we used for this study are not clinical candidates. Nevertheless, our findings strongly support that a therapeutically feasible pharmacological intervention with ASOs to transiently suppress PTBP1 can facilitate the activation of quiescent neurogenic niches to replace neurons within the aged mammalian nervous system.

## Legends of Supplementary Figures

**Supplementary Figure 1: Murine MERFISH gene list design and validation**

**(A)** List of genes used in the initial MERFISH experiments **(Figure 1B-F)** to define cell identity and characterize PTBP1-dependent generation of new neurons. **(B)** UMAPs and spatial maps depicting the expression levels per cell of known markers of distinct cell types in the murine brain organized by neuronal (pink), intermediate (orange) and glial (blue) markers **(C)** Pearson correlation heatmap comparing the major cell type clusters identified by MERFISH (8-week-old mouse) and clusters from the Allen Brain Atlas snRNAseq datasets^88^ (**D)** Histogram with the total number of transcripts detected per cell by snRNA-seq performed by the Allen Brain Atlas^88^ across the 223 genes we measured by MERFISH. The median number of transcripts detected per cell is marked in red. (**E**) Same as (**D**) for the number of genes with at least 10 transcripts per single-cell. **(F)** List of additional genes included to complement the initial MERFISH library.

**Supplementary Figure 2: Comparing cell type definition and gene expression between single-nucleus RNA sequencing and MERFISH data.**

**(A)** Illustration of the experimental design intended to compare MERFISH with single nuclear mRNA sequencing (snRNAseq). 1-year-old wildtype mouse brains were dissected into two hemispheres, with the left hemisphere imaged by MERFISH and the right hemisphere submitted for 10X Genomics snRNAseq. **(B)** UMAP representations of the cells from a 1-year-old mouse coronal brain section imaged by MERFISH – reproduced from Figure 1C for comparison **(C)** UMAP representation of the cells from the snRNAseq data. **(D)** A Pearson correlations heatmap comparing the gene expression of the major cell types identified by both methods. **(E)** Histogram with the total number of transcripts detected per cell by snRNA-seq across the 223 genes we measured by MERFISH. The median number of transcripts detected per cell is marked in red. **(F)** Same as (E) for the number of genes with at least 10 transcripts per single-cell. **(G)** Violin plots with the mRNA level per cell for *Ptbp1*, *Gad1*, and *Prox1* detected using MERFISH in dentate gyrus neurons, GABAergic neurons, and Ependymal cells. **(H)** Violin plots with the mRNA level per cell for *Ptbp1*, *Gad1*, and *Prox1* detected using snRNAseq in dentate gyrus neurons, GABAergic neurons, and Ependymal cells.

**Supplementary Figure 3: Validation of MERFISH using sequential single molecule FISH (smFISH)**

**(A)** Illustration of the probes used for single molecule FISH (smFISH) experiments. Each probe is designed to target a complementary mRNA sequence (40-base) and contains one out of two unique barcodes (20-base) per gene. The signal is detected by hybridization with adaptor and readout probes^63^. The smFISH probes were synthesized with an acrydite anchor modification on the 5’ end to allow for a covalent incorporation into a thin acrylamide gel cast on top of the sample after hybridization. This enhances stability allowing for hundreds of cycles of hybridization and imaging. **(B)** Representative images (top) and insets (bottom) of smFISH for *Ascl1* in the dentate gyrus of an 8-week-old mouse. *Ascl1* signal is shown in green for the first hybridization round targeting a subset of probes against *Ascl1* (left), and in red after 75 rounds of hybridization targeting the remaining *Ascl1* probes (right). **(C)** Spatial map quantifying the single-cell expression level of *Gfap* using the first set (top) and the second set (bottom) of *Gfap* probes separated by 75 hybridizations. **(D)** Correlation plot of the number of *Gfap* transcripts detected in each cell by the two sets of orthogonal probes against *Gfap*. **(E,F,G)** Spatial map quantifying the single-cell expression level of *Gfap* **(E)** *Bsn* **(F)** and *Prox1* **(G)** by MERFISH (top) and smFISH (bottom) in 8-week-old mice. **(H)** Correlation plot between the mean number of transcripts detected per cell across the 223 genes using MERFISH vs smFISH.

**Supplementary Figure 4: PTBP1 suppression in neurons has minimal effect on gene expression**

**(A**) Pearson correlation heatmap between saline and Ptbp1-ASO injected mice comparing three different neuronal cell types including the dentate gyrus neurons, CA1 neurons and L3 cortical neurons. **(B,C,D)** Violin plots of *Ptbp1*, *Sox11*, *Tbr1*, *Gfap*, and *Clasp1* mRNA levels per cell within mature dentate gyrus **(B),** CA1 **(C),** or cortical L3 **(D)** neurons in saline (blue) and Ptbp1-ASO injected mice. **(E)** Dot plot representation of the top 10 differentially expressed genes in saline control and PTBP1-ASO injected mice in dentate gyrus (top) CA1 (middle) and Cortical L3 (bottom) neurons.

**Supplementary Figure 5: Comparison of MERFISH with published single nuclear sequencing of the adult neurogenic niches**

**(A)** Pearson correlation heatmap between our MERFISH gene expression profiles and the ones measured in Hochgerner et al^70^ single-nuclear RNA sequencing across the main cell clusters involved in SGZ neurogenesis. **(B,C)** Dot-plot graphs for marker genes: *Gfap, Slc1a2, Ascl1, Vim, Dcx, Sox11, Prox1* in the main clusters in Hochgerner et al^70^ **(B)** and our MERFISH data **(C). (D)** Pearson correlation heatmap between our MERFISH gene expression profiles and the ones measured in Cebrian-Silla ^71^ single-nuclear RNA sequencing across the main cell clusters involve in SVZ neurogenesis. **(E-F)** Dot-plot graphs for marker genes: *Ascl1, Vim, Dcx, Sox11, Mki67, Prox1* in the cell types identified in Cebrian-Silla **(E)** and our MERFISH data **(F).**

**Supplementary Figure 6 -Ptbp1-ASOs Intra-cerebro-ventricular delivery mediates generation of DG Stage 3 cells along the aged dentate gyrus and improves memory.**

**(A)** Quantification of cell number of DG Stage 1, DG Stage 2 and DG Stage 3 in 4-week, 8-week and 1-year-old mice 1 week post saline or Ptbp1-ASO injections. **(B)** Representative images of SVZ Stage 3 cells in 1 year old mouse dentate gyrus 2 week post injection of saline (top) or Ptbp1-ASO (bottom) **(C)** Bar graph plotting the number of cells per section (one section for each animal) of DG Stage 1, DG Stage 2 and DG Stage 3 in 1-year-old mice post injections of saline or Ptbp1-ASO (n□=□4,4; two-way ANOVA, *p□=0.05). **(D)** Barnes maze training plot of 1 and 2 years-old mice analyzed 2□months post ICV injection with saline or Ptbp1-ASO, during 4 days of practice. Data are presented as mean□±□S.E.M. (For 1 year old cohort n=12,13, respectively; For 2 years old cohort; n□=□4,5, respectively, two-way ANOVA, *p□=□0.0062).

**Supplementary Figure 7 – Human MERFISH gene expression**

**(A)** Histogram with the total number of transcripts detected per cell of the ∼1200 genes we measured by MERFISH in human sample. The median number of transcripts detected per cell is marked in red. (**B**) Same as (**A**) for the number of genes with at least 10 transcripts per single-cell. **(C)** *VIM, SOX2, GFAP* and *C1QL3* transcripts per cell depicted in the UMAP of the human SVZ as well as in their original position in the tissue.

## Methods

### Mouse model and procedures

C57BL/6mice were obtained from The Jackson Laboratory. All procedures were conducted in accordance with the guidelines of The University of California San Diego Institutional Animal Care and Use Committee. Housing conditions, including daily dark/light cycle (6:00 to 18:00), ambient temperature (68–79□degrees F) and humidity (30–70%), were maintained. Ptbp1-ASO (500□μg) or PBS were administered (total of 5□μl) by ICV injection into 1-year-or 2-year old mice. All experimental procedures were approved by the Institutional Animal Care and Use Committee of the University of California, San Diego.

### Non-Human Primates (NHP)

Control NHP tissues obtained from Ionis’s tissue bank from studies previously performed at Labcorp Münster, GERMANY and were approved by their Institutional Animal Care and Use Committee. 4 mm coronal sections were freshly collected from left hemisphere of the adult female cynomolgus monkeys weighing ∼2.5 kg. The samples were frozen immediately and stored in −80. 16 µm brain cryosections were cut using a Leica 2800E Frigocut cryostat at −20°C. Sections on coverslips were fixed with 4% PFA for 10 minutes and continue with the immunolabelling protocol described below.

### Human autopsy

Human tissue was obtained from the UCSD ALS tissue repository that was created following HIPAA compliant informed consent procedures approved by Institutional Review Boards (either Benaroya Research Institute, Seattle, WA IRB# 10058 or University of California San Diego, San Diego, CA IRB# 120056). Brain tissues were acquired using a short-postmortem interval acquisition protocol of 5.57 hours. Tissues were immediately dissected in the autopsy suite, placed in labelled cassettes and frozen at −80C. For this study, we evaluated a Caucasian male ALS patient carrying a mutation in c9orf72 who died at 68 years old after 1.75 years of disease onset.

### Antisense Oligonucleotides

Synthesis and purification of all chemically modified oligonucleotides were performed as previously described^46^. The ASOs were 20 nucleotides in length, wherein the central nucleotides were 2’-deoxyribonucleotides and flanked on both 5’ and 3’ sides by five 2’-methoxyethyl (MOE) modified nucleotides. Internucleotide linkages were phosphorothioates interspersed with phosphodiesters, and all cytosine residues were 5’-methylcytosines. An initial screen of 196 ASOs (transfected into 4T1 cells by electroporation) identified top candidates, which were then further screened in mice by ICV injection for in vivo efficacy and minimal toxicity. The sequence of the PTBP1 ASO use in this study is 5’-GTGGAAATATTGCTAGGCAC-3’.

### Mouse tissue Dissection

Mice were intracardially perfused with Sorenson’s solution for 5 minutes and with a 4% PFA in PBS for 15 minutes. The brain was dissected, post-fixed in the same 4% PFA for 2hours and transferred to 30% sucrose in PBS for at least 2 days at 4 C, then embedded in optimal cutting temperature (OCT) compound and snap-frozen in isopentane (2-methylbutane) cooled at – 40°C. 30 µm brain cryosections were cut using a Leica 2800E Frigocut cryostat at −20°C and stored in PBS with 0.2% sodium azide at 4C.

### Immunofluorescence staining and confocal imaging

The free-floating brain sections were washed 3 times with 1X PBS, incubated in blocking solution (0.5% Tween-20, 1.5% BSA in 1X PBS) for 1hour at room temperature (RT), incubated overnight at RT in antibody diluent (0.3% Triton X-100 in 1X PBS) containing the primary antibodies (see Supplementary Table 1), washed 3 times with 1X PBS and incubated with secondary antibody (Jackson Immunoresearch), diluted in 0.3% Triton X-100 in 1X PBS), washed 3 times with 1X PBS and incubated 10 min with DAPI diluted in 1X PBS (Thermo Fisher Scientific, 100 ng/ml). Sections were mounted on Fisher brand Super frost Plus Microscope Slides (Thermo Fisher Scientific) with Prolong Gold antifade reagent (Thermo Fisher Scientific). Confocal microscopy was performed using a Nikon Ti microscope with a C2 confocal camera with a Plan apo lamda 100x oil NA 1.45 objective and Plan apo lamda 60x oil na 1.4 objective.

### Behavioral tests

Barnes maze test: This is a spatial memory test^101^ sensitive to impaired or enhanced hippocampal function. Mice learn to find an escape chamber (19 x 8 x 7 cm) below one of twenty holes (5 cm diameter, 5 cm from perimeter) below an elevated brightly lit and noisy platform (75 cm diameter, elevated 58 cm above floor) using cues placed around the room. Spatial learning and memory were assessed across 4 trials (maximum time was 3 min) and then directly analyzed on the final (5th) probe trial in which the tunnel was removed and the time spent in each quadrant was determined and the percent time spent in the target quadrant (the one originally containing the escape box) was compared with the average percent time in the other three quadrants. This is a direct test of spatial memory as there is no potential for local cues to be used in the mouse’s behavioral decision.

### MERFISH gene selection and Probe Library Design and Construction

We designed a panel of 223 genes (used in Figure 1) to be imaged with MERFISH^66^. We first identified differential gene markers for each of the 82 subpopulations in prior scRNA-seq data^88^ using differential gene expression (DGE) analyses. The list was then filtered for genes that were either not long enough to construct 48 specific target regions (each 30-nucleotide long) or whose expression levels were outside the range of 0.01 to 300 average UMI per cluster, as measured by scRNA-seq. MERFISH assays were then performed with the MERFISH encoding probe set as shown in Supplementary Table 1. We used 223 binary barcodes to encode RNAs and chose 10 unassigned barcodes to serve as blank controls. The encoding probe set we used contained on average 48 encoding probes per gene, with each encoding probe containing three of the four readout sequences assigned to each gene. The target sequences specific for each gene were designed as previously^75^.

We designed an additional library with 40 bp targeting region which included all the 223 genes mentioned above and extend it with another 52 genes selected based on recent snRNAseq data sets on adult neurogenesis both in the hippocampus^70^ and subventricular zone^71^. The selection of the target sequences was designed as previously^75^. These probes were designed to have two unique readouts for each gene to enable two color serial smFISH (Figure S3). After validation, the MERFISH strategy using this encoding library was implemented by pooling together for each hybridization specific set of adaptor/readout sequences targeting defined subset of genes in each hybridization round (see Supplementary Table 1). This strategy was used in subsequent murine experiments (Figures 2-6).

For the experiments on human tissue we generated a 1130 gene library including 357 genes for cell typing selected based on the Allen’s Institute’s single nuclear RNA sequencing (snRNAseq) dataset for human brain, and other genes relevant for neurodegenerative diseases (see Supplementary Table 1). The selection of the target sequence was similar as described above and the full list of encoding probes and adaptor probes are listed in Supplementary Table 1.

### Microscope setup used for MERFISH imaging

Image acquisition was performed using custom-built microscope-microfluidics systems^63,74^. The systems were built either around a Nikon Ti-U microscope body or an ASI microscope body with a Nikon CFI Plan Apo Lambda 60x oil immersion objective. These systems were built using a similar component list as published previously^75^.

All system components were controlled using custom software, based on https://github.com/ZhuangLab/storm-control.

### Probe synthesis

Primary/encoding probes were amplified from the template library described in Supplementary Tables 1. The generation of the encoding probe sets were prepared from oligonucleotide pools, as described previously^63^. Briefly, we first used limited-cycle PCR to amplify the oligo pools (Twist Biosciences) for approximately 20 cycles. Then, the resulting product was purified via column purification and underwent further amplification and conversion to RNA by a high-yield in-vitro transcription reaction using T7 polymerase (NEB, E2040S). Subsequently, RNA products were converted to single strand DNA with Maxima Reverse Transcriptase enzyme (Thermo Scientific, EP0751), and then purified via alkaline hydrolysis (to remove RNA templates) and DNA oligo purification kits (Zymo Research D4060). The reverse transcription, for the probes used across most experiments (Figure 2-7), was performed with a primer which introduced an acrydite anchor at the 5’ end of the probes. This allowed for the incorporation of the probes into a protective acrylamide gel post sample hybridization.

All primers were purchased from Integrated DNA Technologies (IDT).

### Readout and adaptor probe preparation

All readout and adaptor probes were ordered from IDT (see Supplementary Table 1) and were diluted directly from this stock.

### MERFISH sample preparation

MERFISH samples in Figure 1 were prepared similarly to as described in^66^. Briefly, frozen brains in OCT block were sectioned at −16°C on a cryostat (Leica CM3050S). A series of 16 μm coronal or sagittal slices were cut. MERFISH measurements of 223 genes with 10 non-targeting blank controls was done as previously described, using the encoding sequences (Supplementary Table 1) and published readout probes^76^. Briefly, 16-μm-thick tissue sections were mounted on 40 mm #1.5 coverslips that were silanized and poly-L-lysine coated and subsequently pre-cleared by immersing into 70% (vol/vol) ethanol. Then the tissues were preincubated with hybridization wash buffer (40% (vol/vol) formamide in 2x SSC) for 10 minutes at room temperature. After preincubation, the coverslip was moved to a fresh 60 mm petri dish and added with 50 ul of encoding probe hybridization buffer (2X SSC, 50% (vol/vol) formamide, 10% (wt/vol) dextran sulfate, and a total concentration of 5 uM encoding probes and 1 μM of anchor probe: a 15-nt sequence of alternating dT and thymidine-locked nucleic acid (dT+) with a 5′-acrydite modification (Integrated DNA Technologies). The sample was placed in a humidified 47°C oven for 18 to 24 hours then washed with 40% (vol/vol) formamide in 2X SSC, 0.5% Tween 20 for 30 minutes at room temperature. Samples were post-fixed with 4% (vol/vol) paraformaldehyde in 2X SSC and washed with 2X SSC with murine RNase inhibitor for five minutes. To anchor the RNAs in place, the encoding-probe–hybridized samples were first washed with a de-gassed 4% polyacrylamide solution, consisting of 4% (vol/vol) of 19:1 acrylamide/bis-acrylamide (BioRad, 1610144), 60 mM Tris⋅HCl pH 8 (ThermoFisher, AM9856), 0.3 M NaCl (ThermoFisher, AM9759) supplemented with the polymerizing agents ammonium persulfate (Sigma, A3678) and TEMED (Sigma, T9281) at final concentrations of 0.01% (wt/vol) and 0.05% (vol/vol), respectively. The coverslips were then washed again for 2 min with the same 4% polyacrylamide (PA) gel solution supplemented with the polymerizing agents ammonium persulfate (Sigma, A3678) and TEMED (Sigma, T9281) at final concentrations of 0.03% (wt/vol) and 0.15% (vol/vol), respectively. The gel was then allowed to cast for 5 h at room temperature. The coverslip and the glass plate were then gently separated, and the PA film was incubated with a digestion buffer consisting of 50 mM Tris⋅HCl pH 8, 1 mM EDTA, and 0.5% (vol/vol) Triton X-100 in nuclease-free water and 1% (vol/vol) proteinase K (New England Biolabs, P8107S). The sample was digested in this buffer for ≥36h in a humidified, 37°C incubator and then washed with 2xSSC three times. The samples were finally stained with an Alexa 488-conjugated readout oligo (Integrated DNA Technologies) quantifying polyA and DAPI solution at 1 μg/ml.

MERFISH samples in Figures 2-7 were prepared similarly to as described above with small modification. Briefly, 16-μm-thick tissue sections were pre-cleared by immersing into 70% (vol/vol) ethanol and then 5% SDS in PBS for 4 minutes. Then the tissues were preincubated with hybridization wash buffer (40% (vol/vol) formamide in 2x SSC) for ten minutes at room temperature. After preincubation, the coverslip was moved to a fresh 60 mm petri dish and added with 50 ul of encoding probe hybridization buffer (2X SSC, 50% (vol/vol) formamide, 10% (wt/vol) dextran sulfate, and a total concentration of 1 μg/μl encoding probes. The sample was placed in a humidified 47°C oven for 18 to 24 hours then washed with 40% (vol/vol) formamide in 2X SSC, 0.1% Tween 20 for 30 minutes at room temperature. To anchor the RNAs in place, the encoding-probe– hybridized samples were embedded in 4% polyacrylamide gel for 6h, post-fixed with 4% (vol/vol) paraformaldehyde in 2X SSC and washed with 2X SSC. Human MERFISH sample preparation was also photo-bleached for 6h using a MERSCOPE Photobleacher (cat #10100003, Vizgen). See Supplementary Table 1 for details of encoding library, adaptor and readout used.

### Sequential hybridization for sequential or combinatorial FISH imaging on the microscope

Each round of hybridization consisted of the following general steps: 1) Flow in the hybridization buffer with a set of oligonucleotide adaptor probes specific to each round, as described below, 2) Incubate for 30-90 minutes at room temperature, 3) Flow wash buffer, 4) Incubate for ∼200 s, 5) Flow in the hybridization buffer with readout probes, 6) Incubate for 30 minutes at room temperature, 7) Flow wash buffer, 8) Incubate for ∼200 s and 9) Flow imaging buffer.Imaging buffer was prepared as described previously^75^..

The hybridization buffer and wash buffer were made up of 35% and 30% formamide in 2x SSC, respectively, with the hybridization buffer also containing 0.01% v/v Triton-X. The hybridization buffer was kept separately for each hybridization round and contained two (for human) or three (for mouse) sets of readout probes.

Before the next round of readout probe or adaptor probe hybridization, fluorescent signals from the readout probes in the current round were removed using TCEP cleavage (Figure 1) or by flowing 100% formamide and then 2X SCC (Figures 2-7).

### Image acquisition

For each experiment, we always selected for imaging at least SVZ and hippocampus brain regions in mice and the SVZ in human. After each round of hybridization, we acquired z stack images of each FOV in 3 or 4 colors: 647 nm and 750 nm illumination (or 560 nm, 647 nm, and 750) were used to acquire FISH signal, and 405 nm illumination was used to image DAPI used for image registration. Images were acquired in all channels before the stage was moved in z and images were acquired at a rate of 10-20 Hz.

### For Figure 1 MEFISH processing was achieved using MERlin

Individual RNA molecules were decoded using MERlin v0.6.1 as previously described^65^. Images were aligned across hybridization rounds by maximizing phase cross-correlation on the fiducial bead channel to adjust for drift in the position of the stage from round to round. Background was reduced by applying a high-pass filter and decoding was then performed per-pixel. For each pixel, a vector was constructed for the 22 brightness values from each of the 22 rounds of imaging. These vectors were then L2 normalized and their Euclidean distances to each of the L2 normalized barcodes from MERFISH codebook was calculated. Pixels were assigned to the gene whose barcode they were closest to, unless the closest distance was greater than 0.512, in which case the pixel was not assigned a gene. Adjacent pixels assigned to the same gene were combined into a single RNA molecule. Molecules were filtered to remove potential false positives by comparing the mean brightness, pixel size, and distance to the closest barcode of molecules assigned to blank barcodes to those assigned to genes to achieve an estimated misidentification rate of 5%. The exact position of each molecule was calculated as the median position of all pixels consisting of the molecule. In order to fix the DAPI segmentation quality further, we performed a flat field correction to the dax images. This process fixes the gradient that is present in the DAPI staining dax images so that we don’t get that gradient in our final spatial images.

Cellpose v1.0.2^86^ was used to perform image segmentation to determine the boundaries of cells and nuclei. The nuclei boundaries were determined by running Cellpose with the ‘nuclei’ model using default parameters on the DAPI stain channel of the pre-hybridization images. Cytoplasm boundaries were segmented with the ‘cyto’ model and default parameters using the polyT stain channel. RNA molecules identified by MERlin were assigned to cells and nuclei by applying these segmentation masks to the positions of the molecules. Any segmented cells that did not have any barcodes assigned were removed before constructing the cell-by-gene matrix.

### MERFISH analysis for Figures 2-7

The image analysis pipeline used for the subsequent MERFISH experiments followed a similar principle as for MERLIN but operated on fitted spots rather than on pixels.

### Cell clustering analysis of MERFISH

Combining cell segmentation with the MERFISH single-molecule detection, we constructed cell-by-gene matrices. These matrices were used in combination with the Scanpy version 1.8 package to further analyze the MERFISH data^110^ using python version 3.9 for processing MERFISH data. Count normalization, Principal Component Analysis (PCA), neighborhood graph construction, and Uniform Manifold Approximation and Projection (UMAP) were performed with SCANPY’s default parameters. We performed Leiden clustering using multiple resolution parameters. The top 20 differential genes identified by the rank_gene_groups function were used to annotate each cluster.

### DNA synthesis labelling

To permanently mark cells undergoing DNA replication, 100 μL of 200mg/ml 5-ethynyl-2’-deoxyuridine (EdU; E10187, ThermoFisher Scientific) was administered daily for 2 weeks by intra perennial injections into 4 week, and 1 years old mice injected either with saline or PTBP1-ASO. Brains were collected 2 weeks post Edu labeling (1 month post PTBP1-ASO delivery). The EdU incorporated in the genome of replicating cells was developed using click-chemistry. The following steps were added in the MERFISH protocol described above after gel casting: 1) washing and blocking with 3%BSA in PBS three times for 5 minutes each, 2) incubating brain sections with Click-iT Plus reaction cocktail (made following the manual on the Click-iT Plus EdU Imaging Kits, C10637, ThermoFisher Scientific) for 30 minutes protected from light at room temperature, 3) washing with PBS three times for 5 minutes each. Then the sample continued with the MERFISH protocol.

### Single-nucleus RNA-seq data generation

Single nucleus RNA-seq was performed by the Center for Epigenomics at UC San Diego using the Droplet-based Chromium Single-Cell 3’ solution (10x Genomics, v3 chemistry)^111^. Briefly, 30 mg brain tissues were resuspended in 500 µL of nuclei permeabilization buffer (0.1% Triton-X-100 (Sigma-Aldrich, T8787), 1X protease inhibitor, 1 mM DTT, and 0.2 U/µL RNase inhibitor (Promega, N211B), 2% BSA (Sigma-Aldrich, SRE0036) in PBS). Sample was incubated on a rotator for 5 min at 4°C and then centrifuged at 500 rcf for 5 min (4°C, run speed 3/3). Supernatant was removed and pellet was resuspended in 400 µL of sort buffer (1 mM EDTA 0.2 U/µL RNase inhibitor (Promega, N211B), 2% BSA (Sigma-Aldrich, SRE0036) in PBS) and stained with DRAQ7 (1:100; Cell Signaling, 7406). 75,000 nuclei were sorted using a SH800 sorter (Sony) into 50 µL of collection buffer consisting of 1 U/µL RNase inhibitor in 5% BSA; the FACS gating strategy sorted based on particle size and DRAQ7 fluorescence. Sorted nuclei were then centrifuged at 1000 rcf for 15 min (4°C, run speed 3/3) and supernatant was removed. Nuclei were resuspended in 35 µL of reaction buffer (0.2 U/µL RNase inhibitor (Promega, N211B), 2% BSA (Sigma-Aldrich, SRE0036) in PBS) and counted on a hemocytometer. 12,000 nuclei were loaded onto a Chromium Controller (10x Genomics). Libraries were generated using the Chromium Single-Cell 3′ Library Construction Kit v3 (10x Genomics, 1000075) with the Chromium Single-Cell B Chip Kit (10x Genomics, 1000153) and the Chromium i7 Multiplex Kit for sample indexing (10x Genomics, 120262) according to manufacturer specifications. CDNA was amplified for 12 PCR cycles. SPRISelect reagent (Beckman Coulter, B23319) was used for size selection and clean-up steps. Final library concentration was assessed by Qubit dsDNA HS Assay Kit (Thermo-Fischer Scientific) and fragment size was checked using Tapestation High Sensitivity D1000 (Agilent) to ensure that fragment sizes were distributed normally about 500 bp. Libraries were sequenced using the NextSeq500 and a NovaSeq6000 (Illumina) with these read lengths: 28 + 8 + 91 (Read1 + Index1 + Read2).

### Integration of single-cell RNAseq and MERFISH

To integrate the scRNA-seq and MERFISH datasets, we first subset both datasets to only genes interrogated by both modalities. We then utilized SCANPY’s implementation of Harmony to project both the scRNA-seq and MERFISH datasets into a shared PCA space^112^. The dimensionality of the joint embedding was further reduced using UMAP (min_dist=0.3) to visualize the space in two dimensions using a k-nearest neighbor graph constructed with a k of 30 and the Pearson correlation distance metric to match the parameters used by the Harmony authors. To impute a complete expression profile and cell type for each MERFISH profile, we assigned the expression profile and cell type of the closest scRNA-seq cell in the Harmony PCA space using the Euclidean distance metric.

### Statistics and graphs

All statistical analyses and graphs were performed using GraphPad Prism v8.0. For two-group analysis, Student’s t-test or the Mann-Whitney test was used, as determined by a normality test. All experiments include at least 3 biologically independent repeats; significance was set at *p<0.05, **p<0.01, ***p<0.001. Statistical significance was set at p< 0.05, and data are presented as mean ± SEM.

## Acknowledgements

This work was supported by research grants from the NIH (R01-NS27036 from to D.W.C. and B.B. and DP5-OD030878 to B.B.). R.M. was supported by postdoctoral fellowships from the Hereditary Disease Foundation and the Huntington Disease Society of America and by a research grant from the Dan Lewis Foundation. We thank the Center for Epigenomics at UC San Diego for performing the single nucleus RNA-seq experiments. Work at the Center for Epigenomics was supported in part by the UC San Diego School of Medicine: IGM Genomics Center, UC San Diego. This publication includes data generated at the UC San Diego IGM Genomics Center utilizing an Illumina NovaSeq 6000 that was purchased with funding from the NIH (S10 OD026929).

## Author Contributions

R.M., C.C.-M., S.V.-S., D.W.C. and B.B conceived the study. R.M., C.C.-M., S.V.-S., C.K., K.J., Q.Z., D.W.C. and B.B designed the study. R.M., C.C.-M., S.V.-S., C.K., K.J., K.M., S.M., J.C., M.M.-D.,C.H.,P.J.-N., Q.Z., J.R and B.B. performed the experiments. R.M., C.C.-M., S.V.-S., C.K., K.J., S.Moo, S.Mog, A.M, D.W.C., and B.B. analyzed the data. R.M., C.C.-M., S.V.-S., C.K., Q.Z., A.G., C.F.B., D.W.C. and B.B. wrote the manuscript. All authors discussed the results and commented on the manuscript.

