## Supplemental Figures for "Re-activation of neurogenic niches in aging brain"

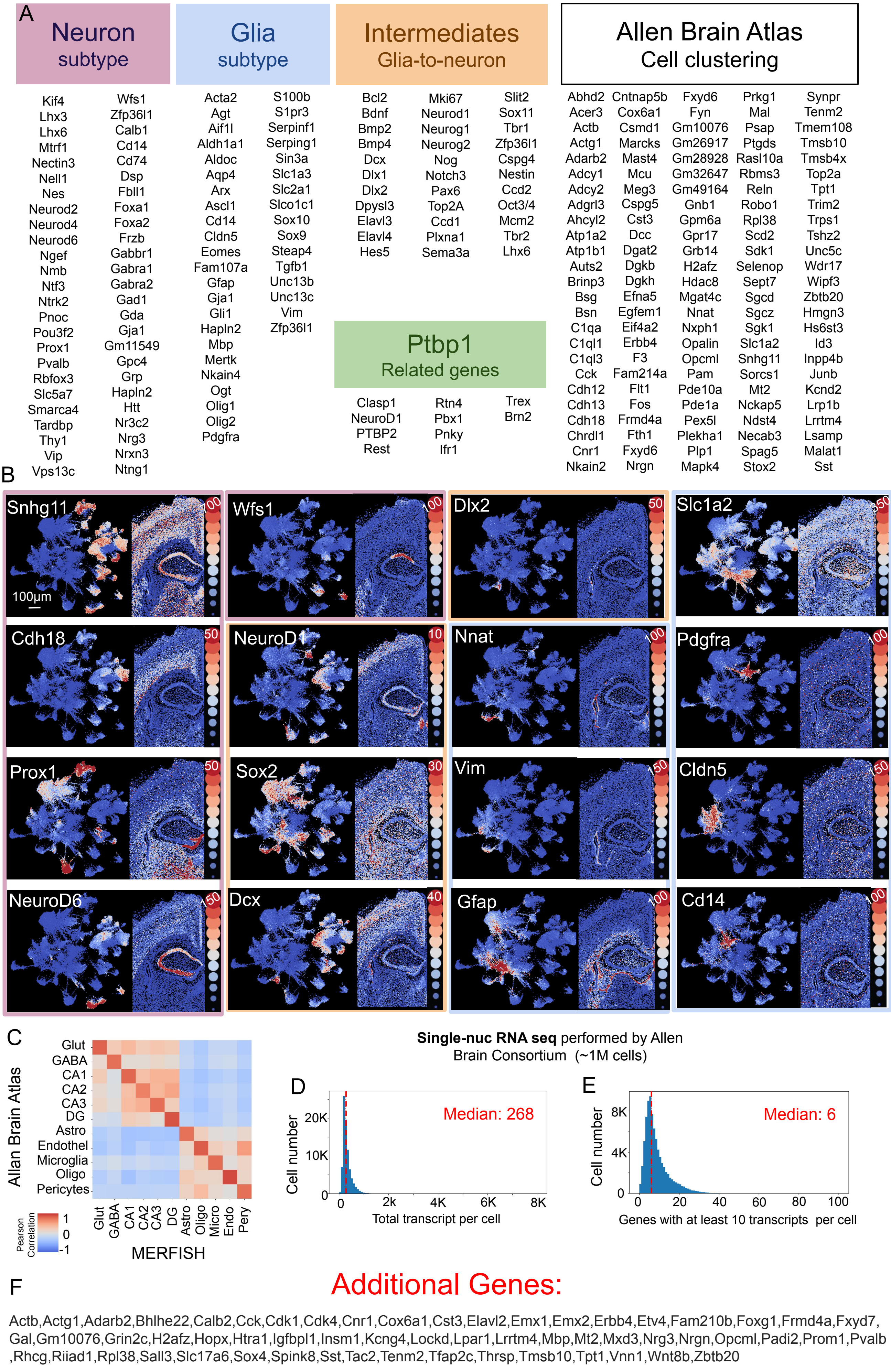


Maimon et al, Supplementary Figure 1: Murine MERFISH genes list construction and validation

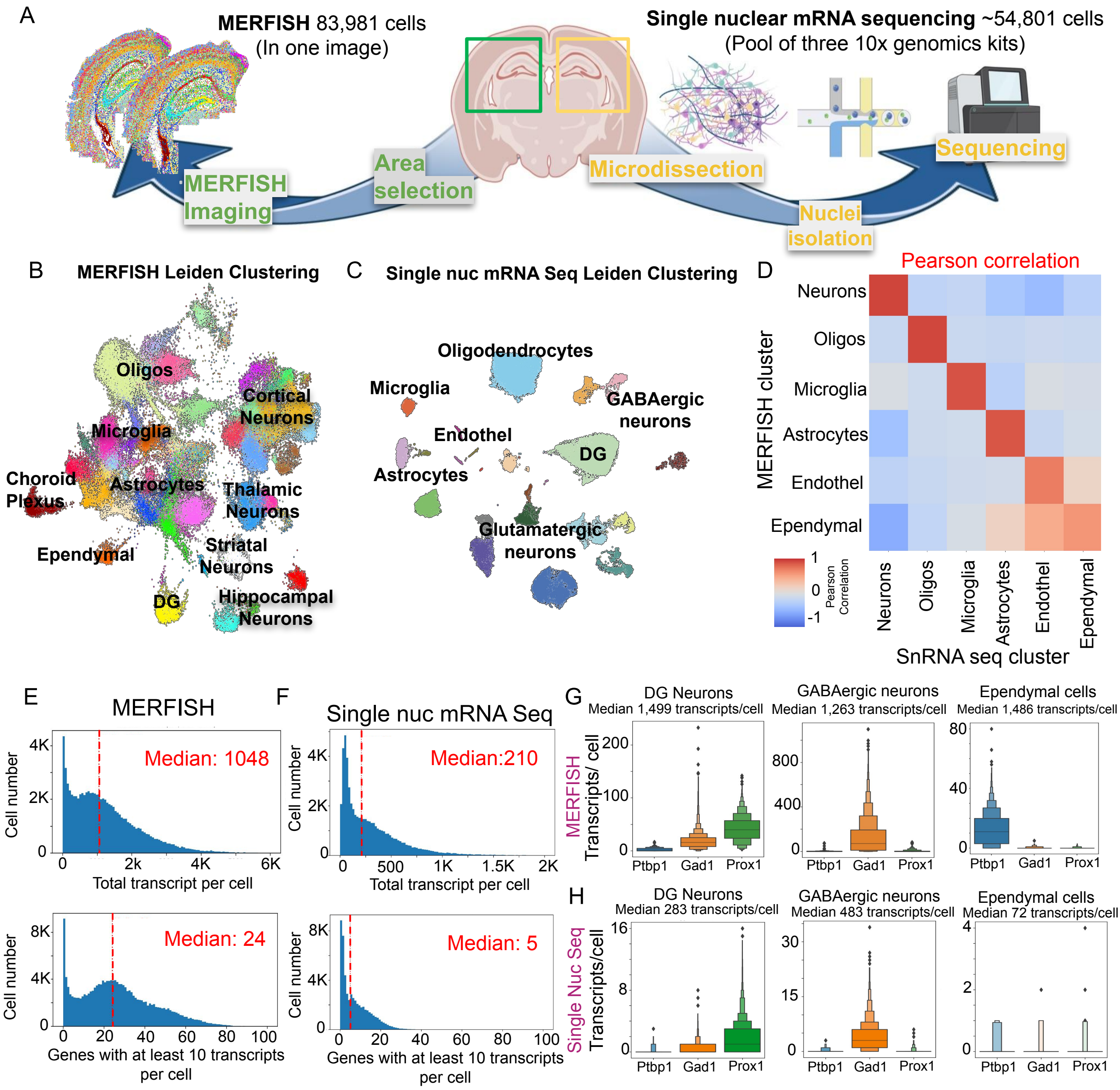


Maimon et al, Supplementary Figure 2 - Comparing cell type definition and gene expression between single-nucleus RNA sequencing and MERFISH data

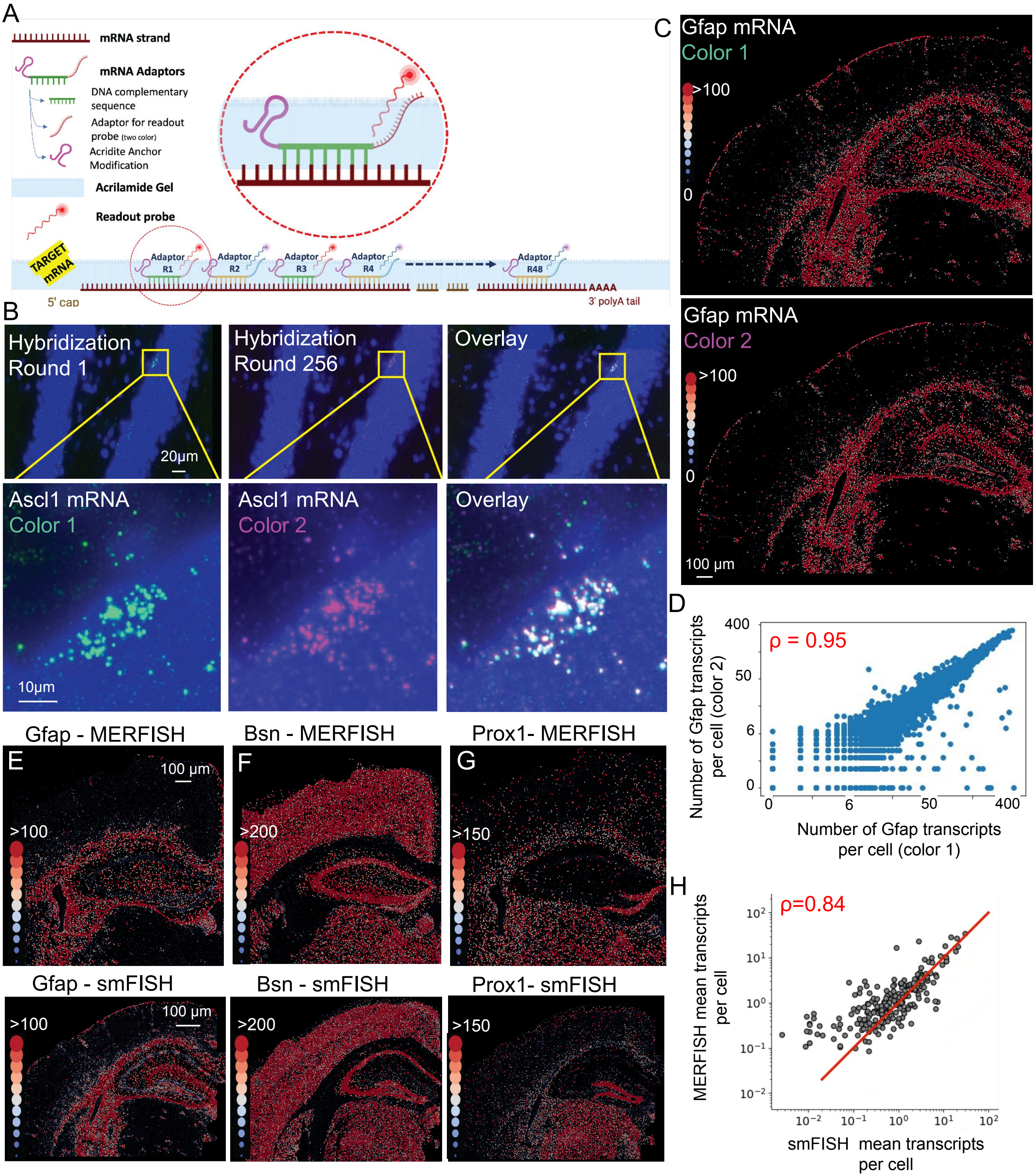


Maimon et al, Supplementary Figure 3: MERFISH validation with serial smFISH

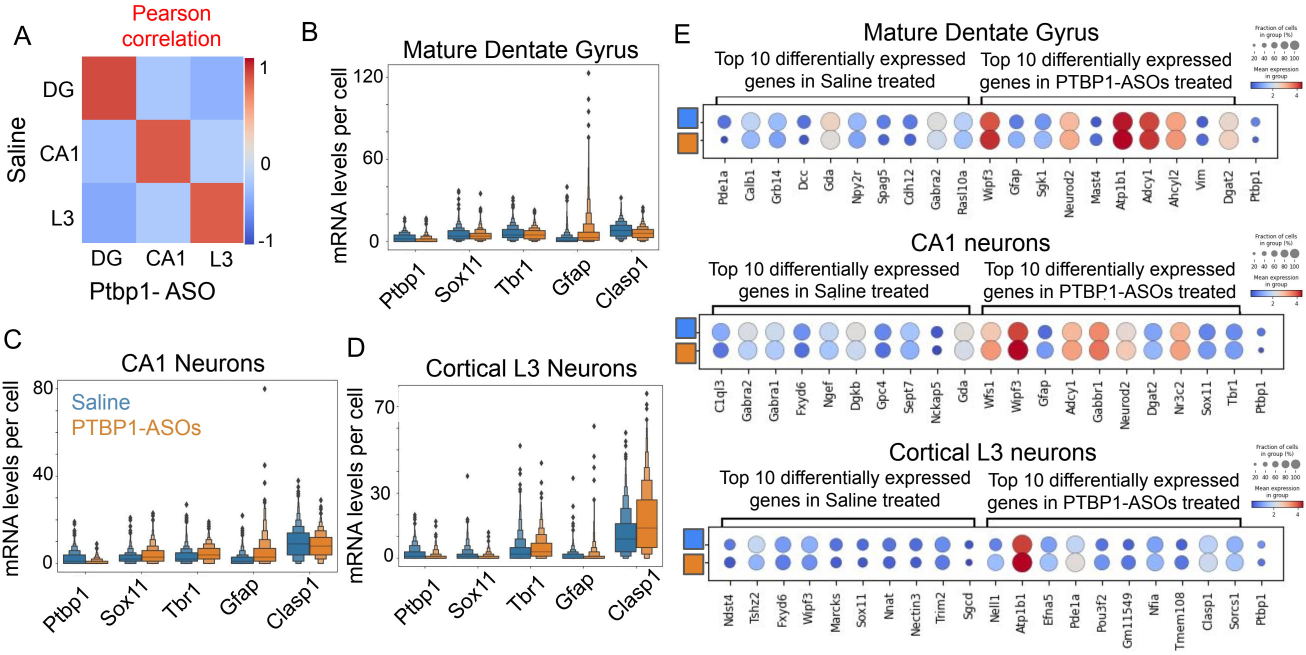


Maimon et al, Supplementary Figure 4: PTBP1 suppression in neurons has minimal effect on gene expression

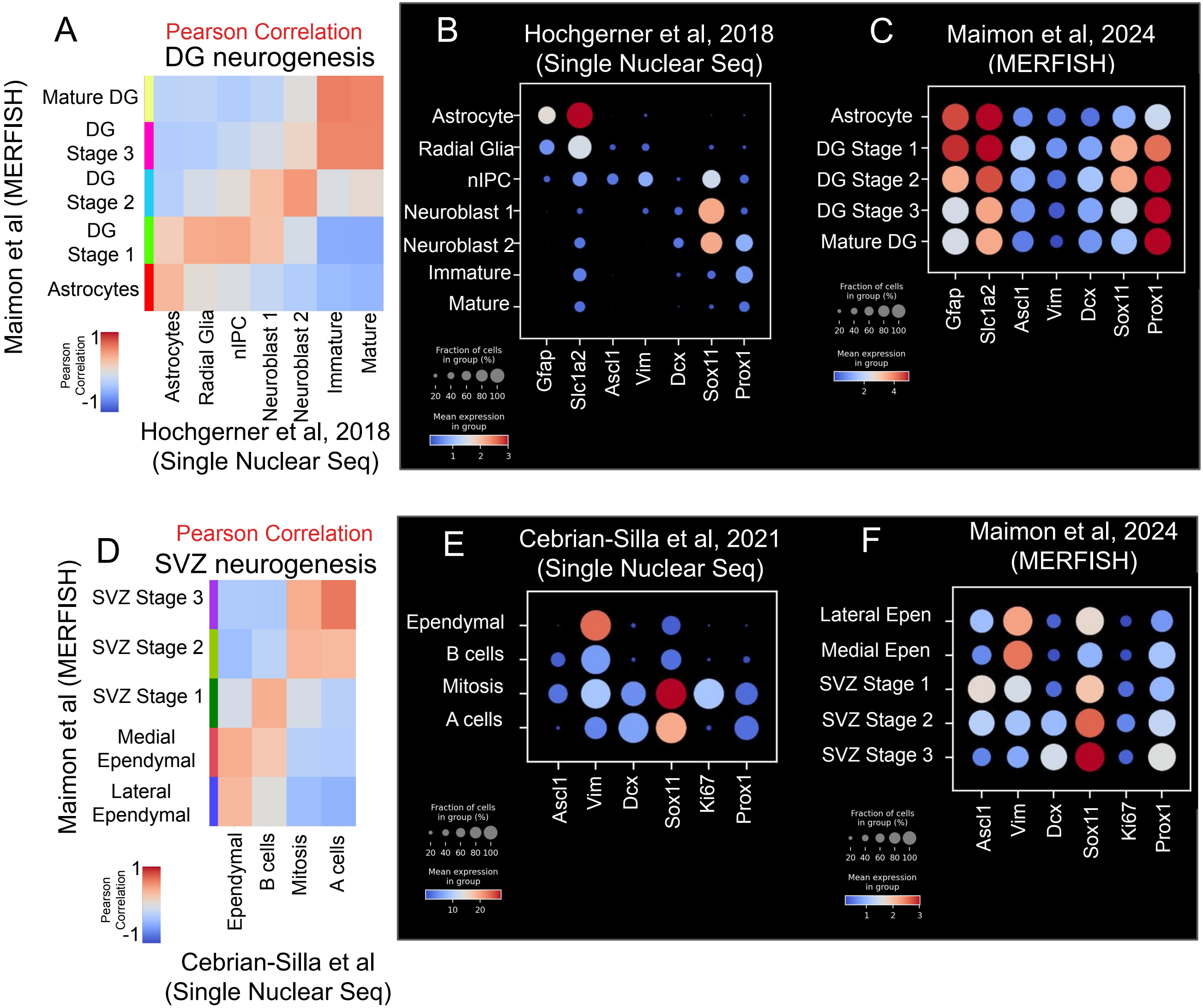


Maimon et al, Supplementary Figure 5: Comparison of dentate gyrus and SVZ single nuclear RNA sequencing with MERFISH

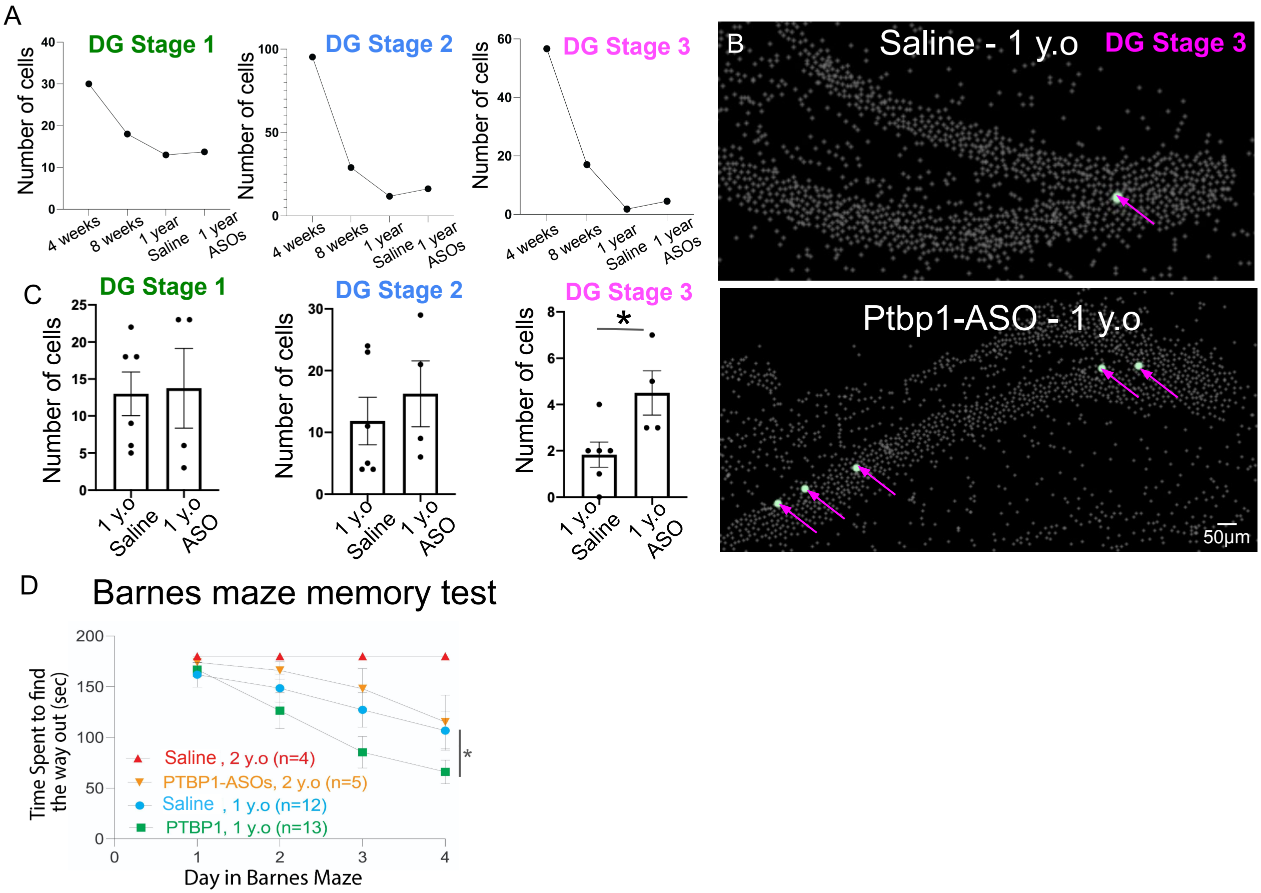


Maimon et al, Supplementary Figure 6: PTBP1-ASOs Intra-cerebro-ventricular delivery mediate generation of DG Stage 3 cells along the aged dentate gyrus and improves memory.

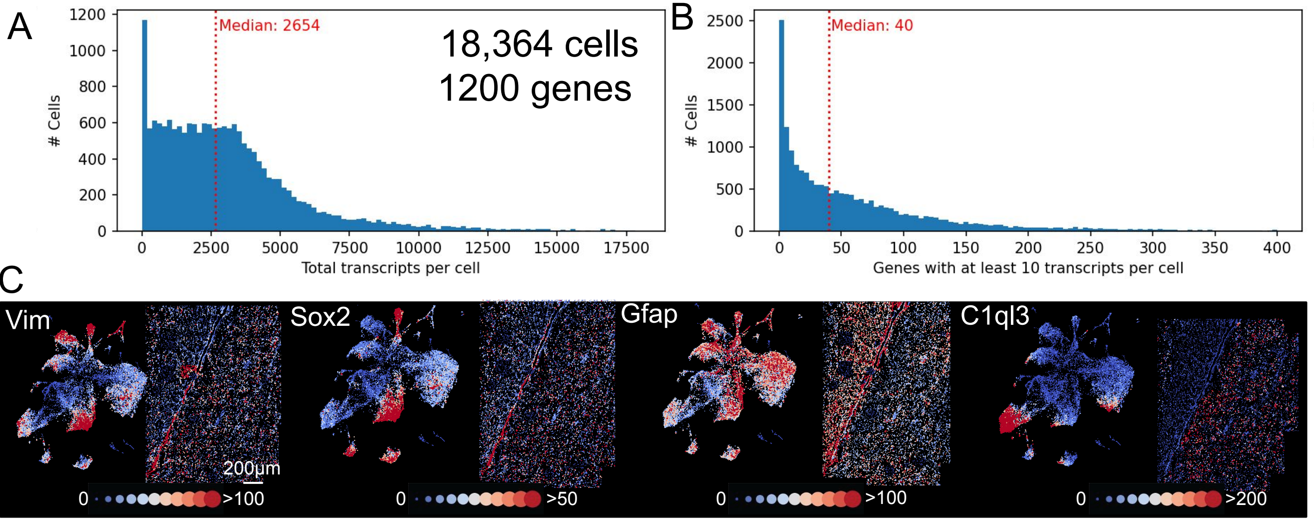


Maimon et al, Supplementary Figure 7: Human MERFISH quality control
